# Transferability of stream benthic macroinvertebrate distribution models to drought-related conditions

**DOI:** 10.1101/2024.05.28.596146

**Authors:** Graciela Medina Madariaga, Hong Hanh Nguyen, Jens Kiesel, Kristin Peters, Christian K Feld, Sonja C. Jaehnig, Yusdiel Torres Cambas

**Affiliations:** Leibniz-Institut of Freshwater Ecology and Inland Fisheries and Geography Department, Humboldt-University of Berlin; Department of River Ecology and Conservation, Senckenberg Research Institute and Natural History Museum Frankfurt, Frankfurt, Germany; Institute for Natural Resource Conservation, Christian-Albrechts-University Kiel, Kiel, Germany; Faculty of Biology, Department of Aquatic Ecology and Centre for Water and Environmental Research, University of Duisburg-Essen, Essen, Germany; Leibniz-Institut of Freshwater Ecology and Inland Fisheries

**Keywords:** Ecological niche modelling, extreme event, climate change, low-flow, extreme flow, prevalence, independent evaluation

## Abstract

Freshwater ecosystems increasingly face pressures from climate change induced extreme events, like droughts, posing significant threats to biodiversity. While Species Distribution Models (SDMs) serve as vital tools for predicting species responses to environmental shifts, their transferability to novel environmental conditions, especially during and after drought remains poorly understood. In this study, we delve into the transferability of SDMs for freshwater macroinvertebrates from drought-free to drought-influenced conditions. We examine how sensitive the transferability is to traits such as tolerance scores according to their distribution along longitudinal gradients, as well as the used modelling method. We constructed and validated SDMs for freshwater macroinvertebrates in a central German catchment under drought-free conditions using four different algorithms (Generalised linear models; GLM, Spatial Stream Networks; SSN, Random Forests; RF, and Maximum Entropy; MaxEnt). We then projected these models to environmental conditions influenced by drought, and obtained their transferability by computing the difference in accuracy when predicting under drought-free and drought-influenced conditions (AUCgap). Our findings reveal a marked reduction in SDM accuracy under drought conditions, illustrating the challenges of accurately predicting species distributions into novel environmental conditions. Our results show a slightly better transferability when using SSN and RF. In addition, we observed that SDM transferability can be influenced by species tolerance, with sensitive and tolerant species presenting higher AUCgap (i.e. lower transferability). Furthermore, we found that when sensitive species were modelled using SSNs, the AUCgap was reduced. Our study underscores the limitations of SDMs in capturing species responses to drought and advocates for integrating ecologically relevant predictors and modelling methods that account for the stream connectivity to allow for robust predictions. These considerations could enhance the ability of SDMs to effectively estimate the impacts of extreme events on freshwater biodiversity.

## 1. Introduction

Freshwater ecosystems host an immense biodiversity and provide essential ecosystem services, yet at the same time are particularly vulnerable to the effects of climate change (Markovic et al., 2017; Comte & Olden 2017). One expected consequence of climate change is the increase in the frequency, magnitude and duration of extreme weather events (e.g., heatwaves, droughts, storms; hereafter called extreme events). Extreme extreme events related to flow conditions in streams and rivers manifest in the form of extended droughts (Messager et al. 2021) and increased number of floods (Kundzewicz et al., 2014; Nilsson et al., 2015; Beniston et al., 2007; Meehl & Tebaldi 2004), and represent a growing and alarming threat, further intensifying the already observed declines in freshwater biodiversity (He et al., 2019; Tickner et al., 2020; WWF, 2022). On biodiversity, the effects of extreme flows differ depending on the type of event, with low flows being associated with more significant decreases in species richness than high flows (Sabater et al., 2023). Droughts can pose adverse effects on fish migration, due to habitat loss (Röpke et al., 2017), and high flows following droughts can disrupt water quality and lead to increased mortality (King et al., 2012). For freshwater macroinvertebrates, droughts can benefit tolerant and invasive species in river ecosystems (Daufresne et al., 2007), and negatively impact the richness and abundance of mayfly, stonefly, and caddisfly (EPT) taxa (Herbst et al., 2019). Indeed, the effects of droughts on freshwater macroinvertebrate biodiversity (Hill et al., 2019) have been estimated to be twice as strong as the effects of a gradual warming (Sabater et al., 2023).

The ability to predict changes in species distributions under climate change scenarios is essential to better tailor management and conservation strategies (Guisan et al., 2013). To date, the distribution of macroinvertebrates has mostly been associated with gradual changes in climatic variables, such as those associated with temperature and precipitation (Domisch et al., 2013; Shah et al., 2014). However, macroinvertebrates are highly sensitive to abrupt changes in environmental conditions (Bertoncin et al., 2019), such as those caused by droughts (Aspin et al., 2019a). Considering the forecasted rise in the frequency and severity of extreme events anticipated in the forthcoming years, predicting how extreme flows might shape species distributions becomes of pressing importance, to better safeguard freshwater biodiversity. Within this scope, Species Distribution Models (SDMs) are a frequently-used tool in ecology for investigating and predicting species distributions. Their versatility and comparatively low data intensity as contrasted with other modelling methods (Peterson et al., 2015), have made them valuable in ecological research.

Nonetheless, SDMs have been faced with the issue of inaccurate predictions when projected to new environments (Eger et al., 2017, Qiao et al., 2019). The challenges posed by extreme flows, characterised by sudden and drastic environmental changes, further accentuate the complexities associated with SDM transferability. In the case of droughts, effects in rivers and streams can translate to habitat loss and a decrease of organic matter accompanied with changes in the water chemistry (Bond et al., 2008). These modifications represent different conditions to those found before drought and their consequences on biota can persist even after reestablishment of flow (Calapez et al., 2014; Ruiz et al., 2022), potentially affecting SDM predictions and questioning the robustness of models and their ability to adequately reflect such abrupt environmental shifts. The ability to successfully transfer a model has become a priority to support informed decision-making under the rapidly changing global environmental landscape (Yates et al, 2018), especially in the context of climate change (Werkowska et al., 2017). Therefore, there is a growing need to evaluate the performance of models under climatic shifts associated with the increase of extreme flows such as droughts.

Beyond fundamental aspects such as variable selection and sample size, species traits play a crucial role in determining the transferability of SDMs to novel conditions (Werkowska et al., 2017). Studies on birds have established associations between model transferability and the longevity of the species modelled (Rousseau & Betts, 2022). In freshwater ecosystems, macroinvertebrate Joint SDMs (JSDMs), which are used to predict multiple species simultaneously, have suggested that specific traits might play a role in the distribution of species (Elo et al., 2021), potentially having implications in their transferability, as shown in research on bryophytes (Collart et al., 2023). Moreover, SDMs for tolerant species have been associated with lower accuracies (Hernandez et al., 2006), and species’ tolerance to environmental conditions might mitigate or intensify their responses to exposure to extreme flows (Chessman et al., 2015). Therefore, traits related to the tolerance of a species might affect how much the accuracy of SDMs can be extrapolated to scenarios under the influence of drought.

In addition, the method employed to develop SDMs exerts a substantial influence on their prediction accuracy. Two main categories of SDM methods can be identified and are more frequently used in ecological studies (Miller, 2010): (1) traditional regression methods such as generalised linear models (GLMs), which model the probability of a species’ distribution as a linear combination of the predictor variables, and (2) machine-learning methods, like Random Forests (RFs) and maximum entropy models (MaxEnt), etc., which are usually “data-driven” and can fit complex relationships. In terms of transferability, extrapolating models trained and evaluated on data from one geographical area may experience accuracy losses ranging from up to 40% when projected to different geographical areas (Gies et al., 2015) and also depending on the method used (Charney et al. 2021). Furthermore, accurate estimation of species distribution in riverine ecosystems is also influenced by the connectivity within the stream network (Fortuna et al., 2006; Altermatt et al., 2013; Baldan et al., 2021). In the context of scenarios associated with drought, the distance to perennial sources and drought refuges may significantly influence species distribution after these events have occurred (Boulton et al., 2003; Van Looy et al., 2019), and this might differ between widely distributed species and species with limited distributions. Defining modelling approaches for SDMs, in particular incorporating methods that account for stream connectivity, may provide a valuable advantage in improving the predicted species distributions related to drought.

Here, we analyse the accuracy of species distribution models (SDMs) in conditions with and without the influence of drought. We explore how SDMs behave according to species traits, particularly the species’ tolerance to environmental conditions (i.e., tolerance score; Graf et al., 2018), and the modelling method used. We also examine the impact of accounting for connectivity through the inclusion of Spatial Stream Networks (SSN) as a modelling approach (Ver Hoef et al., 2006). We aim to answer four main questions: 1) Are SDMs from drought-free conditions transferable to drought-influenced conditions? 2) Do SDMs based on specific tolerance scores yield higher transferability? 3) Are there modelling methods (RF, Maxent, GLM, SSN) that are more transferable? And 4) Does any modelling method transfer better to drought-influenced conditions for certain tolerance scores? We aim to contribute valuable insights into the strengths and limitations of SDMs in predicting species distributions, particularly in the aftermath of drought in river ecosystems.

## 2. Methods

### 2.1. Study area

The Kinzig river is located in the central lower-mountainous ecoregion of Germany (Fig. 1), encompassing a catchment area of approximately 1058 km². The predominant land use is characterised by extensive sylviculture (42%), open land (27%), and cropland (20%). In line with the broader trend of climate warming across Central Europe (IPCC, 2023), the region has witnessed notable instances of prolonged drought (Haase et al., 2019; e.g., most extreme low flows in summer 2016). These climatic shifts have resulted in observable changes in the composition of the macroinvertebrate community, partially attributed to their responses to drought events (Nguyen et al., 2023, 2024).

**Fig 1.**
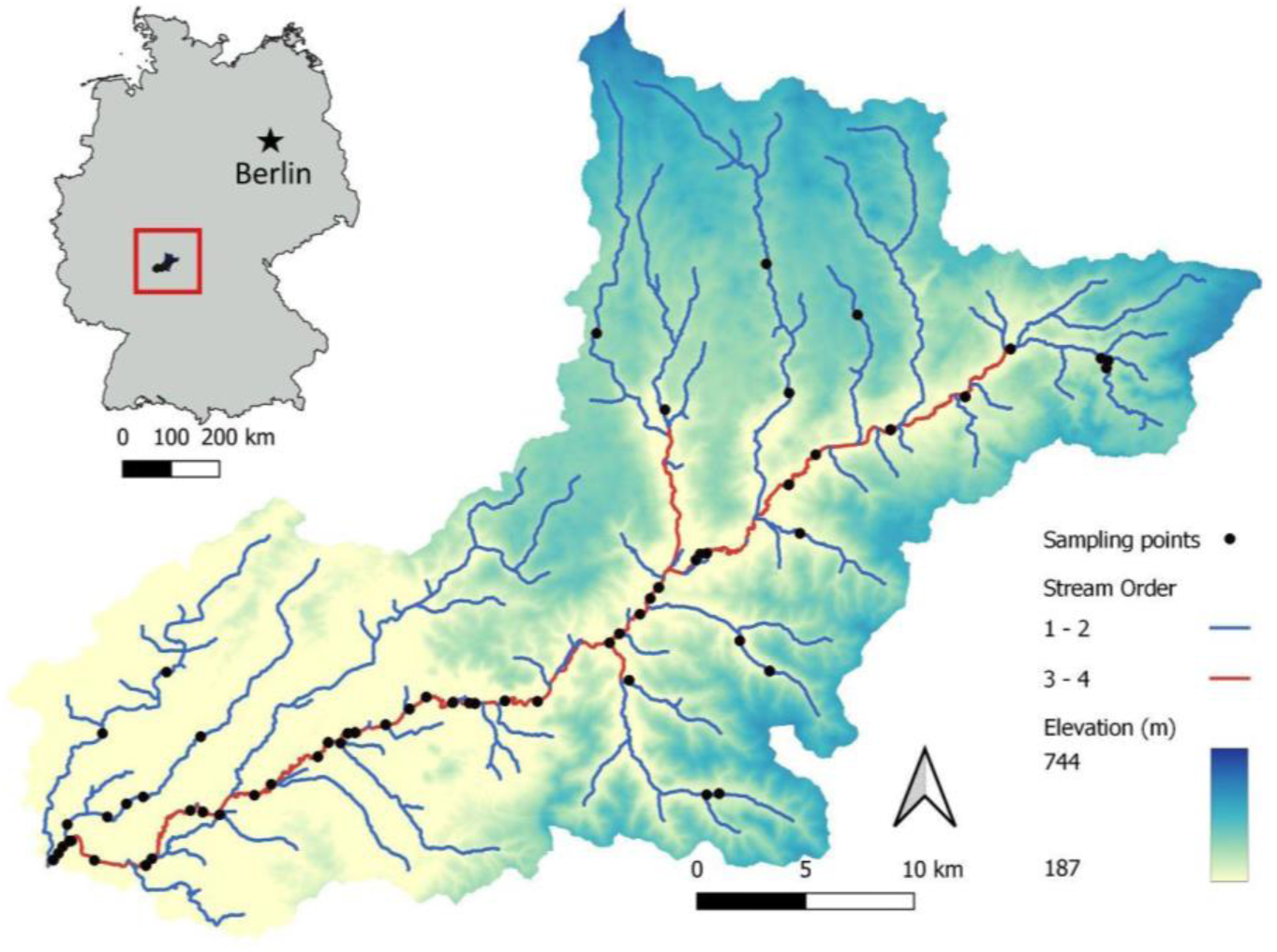
Map of the Kinzig catchment located in central Germany and sampling points.

### 2.2. Biological data

Macroinvertebrate presence and absence records were processed from abundance data obtained from biomonitoring performed by regional authorities and the Rhine-Main-Observatory (https://deims.org/9f9ba137-342d-4813-ae58-a60911c3abc1), spanning the years 2007 to 2021. The samples were collected during two seasons, with tributary and low-order streams sampled during spring, and the sites along the main channel of the Kinzig sampled during summer. Macroinvertebrate sampling (Supporting Information 1.1) was performed following the European Water Framework Directive (EU WFD) compliant German national standard sampling protocol (Meier et al. 2006) and taxa were classified according to the operational taxa list (Haase et al. 2006).

### 2.3. Environmental Data

Climate, elevation, land use, and Indices of Hydrological Alteration (IHAs; Richter et al., 1996) data were assembled to construct our SDMs (Table 1). The variable categories included in the selection process were chosen due to their relevance for shaping the distribution of freshwater macroinvertebrate species. For instance, climate has been found a major driver of aquatic macroinvertebrate distribution in freshwater ecosystems and bioclimatic variables have been successfully used to estimate the distribution of macroinvertebrates in streams globally and regionally (Domisch et al., 2013; Kuemmerlen et al., 2014). Elevation can influence habitat availability and microclimatic conditions, consequently exerting a pronounced impact on the distribution patterns of macroinvertebrates (Madsen et al., 2015). Moreover, land use can have a strong impact in freshwater biota and especially freshwater macroinvertebrates (Schürings et al., 2022). Finally, in river ecosystems, IHAs play a significant role in describing macroinvertebrate trends (Olden & Poff, 2003; Kakouei et al., 2018; Irving et al., 2022).

**Table 1.** Variables produced for SDMs. Data sources and processing can be found in the Supporting Information 1.2.

| Variable category | Variables |
| --- | --- |
| Bioclimatic variables | Climatic parameters derived from temperature and precipitation data, calculated for the 12 months before sampling* |
| Elevation | Elevation |
| Land Use | % of upstream coverage of: forest, open land, cropland and urban surfaces |
| IHAs | Variables describing the magnitude, duration, frequency, timing and rate of flow events during the 12 months before sampling* |

\*A detailed list can be found in Table S1

### 2.4. Species Distribution Models

To assess the transferability of SDMs from drought-free conditions to conditions under drought influence we developed SDMs within the R software (R Core Team, 2023). First, we divided the biological data into two independent subsets representing different exposure to drought. The data partitioning was conducted using a distinct cutoff, which denoted a decline in runoff (Nguyen et al., 2024), as indicated by the runoff rate (spanning the 12 months preceding macroinvertebrate samplings; Fig. S1). The first subset, referred to as Period1, was utilised for modelling species’ distributions before drought was identified, encompassing the years 2007 to 2016. The second subset, Period2, was dedicated to evaluate the predicted distributions of species under environmental conditions influenced by drought and covered the years from 2017 to 2021. We retained only taxa present in more than 10% and less than 90% of all sampled points during Period1, due to the effect of prevalence on SDM accuracy (Gies et al., 2021). This resulted in 42 macroinvertebrate taxa (Table S2).

Species Distribution Models (SDMs) were developed for each taxon using Period1. We utilised four methods: Generalised Linear Models (GLM; Friedman et al., 2010), Spatial Stream Networks (SSN; Ver Hoef et al., 2006), Random Forest (RF; Liaw & Wiener 2002), and MaxEnt (Phillips et al., 2004) for model fitting. Presence-absence data were converted to binary data, and therefore we opted to not generate pseudo-absences. For the validation of models, we randomly divided Period1 into two fractions, one for model training (80% of the data) and another one for validation (20% of the data). For our variable selection process, we used ‘corSelect’ on each variable group (bioclim, land use, elevation, and each subgroup of magnitude, duration, frequency, rate and timing of IHAs) to obtain all pairs of highly correlated variables (≥0.7) and further retaining the variable with the lowest VIF, resulting in 33 predictors. Additionally, ‘select07’ was employed to choose influential variables for taxon presence/absence, the number of predictors was selected based on the species’ prevalence in Period1, and lowest AIC values. Selected variables were used for subsequent SDM using the aforementioned modelling methods. Due to stochasticity, we repeated this process five times (Fig. 2; Guisan et al., 2017), creating five models for each species with respective validation metrics (Period1Acc): Area Under the Curve (AUC) and True Skill Statistic (TSS), resulting in 840 models (42 species x 5 models x 4 methods). Valid models were chosen for further analysis based on specific criteria: AUC greater than 0.7, and a TSS exceeding 0.5 (Allouche et al., 2006; Konowalik & Nosol, 2021). Notably, two species (*Dendrocoelum lacteum* and *Anabolia nervosa*) did not produce valid models within Period1 regardless of the method used and were subsequently removed from further analysis. Valid models were employed to predict species distributions influenced by drought. To assess the accuracy of these models following drought events, confusion matrices were generated by comparing predicted versus observed data (Period2), the accuracy of predictions was stored as Period2Acc.

**Fig 2.**
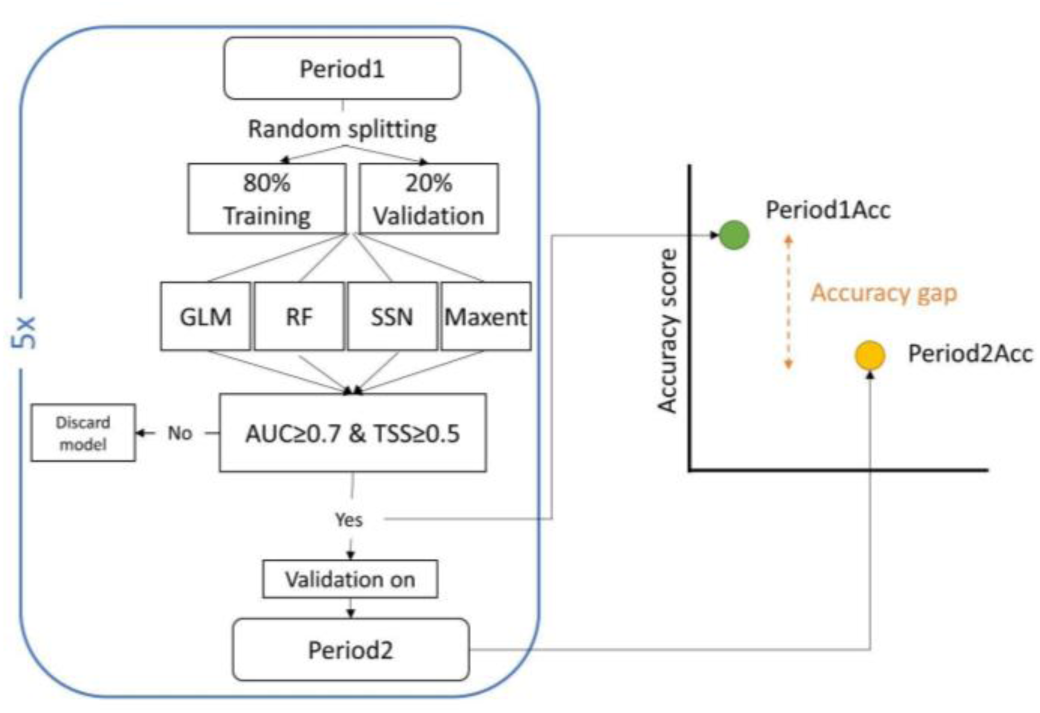
Schematic representation of the accuracy gap determination procedure. The process of training and validating a model on Period1, followed by (external) validation on Period2. The difference in accuracy between Period1 and Period2 defines the accuracy gap (AUCgap).

### 2.5. Statistical analysis

We computed the difference between models’ accuracy gap (AUCgap; Rückauer et al., 2019) obtained during validation on Period1 (Period1Acc) and their accuracy when predicting Period2 presence/absences (Period2Acc). The influence of the method used and the species’ tolerance in the transferability of SDMs were key considerations in our analysis. The species’ tolerance score (Graf et al., 2018), mainly based on species zonation preference (Moog et al., 1995), considers species found in more zones as more tolerant and species with more limited zonation preferences as more sensitive. This metric has shown correlation with the saprobic index and is a simpler metric used to summarise the tolerance of a species to environmental changes, therefore serves as a practical tool for biomonitoring. Additionally, the information on tolerance score is available for a great number of European freshwater macroinvertebrate species. Information on the tolerance score was sourced from freshwaterecology.info (Schmidt-Kloiber & Hering, 2015). Species were categorised as tolerant (tol), moderately tolerant (mtol), moderately sensitive (msen), and sensitive (sen). Species belonging to the category highly sensitive (hsen) were not represented in our dataset.

Wilcoxon tests were used to assess differences between Period1Acc accuracy and Period2Acc accuracy. Additionally, Shapiro-Wilk tests (Shapiro & Wilk, 1965) were conducted to assess the normality of the resulting AUCgap (Fig. S2), and simple linear models were used to investigate the effect of species prevalence in the dataset on AUCgap. Kruskal-wallis (KW) tests were performed to discern whether significant differences were obtained between tolerance scores categories (tol, mtol, msen and sen) and across methods (GLM, SSN, RF, MaxEnt; Table S3). When KW resulted in significant differences across groups, pairwise Dunn post-hoc tests were used to identify significant differences between categories.

Acknowledging the inherent variability in model accuracy linked to the species modelled (Tessarolo et al., 2021; Fig. S3), we employed linear mixed models to describe the accuracy gap with “Species” as a random effect (Formula 1: AUCgap ∼1+(1|Species)), using the package *lme4* (Bates, 2015). The significance of each variable was assessed by incorporating species tolerance score, modelling method used, and their interaction as fixed effects into a full model (Formula 2: AUCgap ∼ Tolerance + Method + Tolerance:Method + (1|Species)). We selected the best model describing the variation in AUCgap based on Akaike Information Criterion (AIC), R² conditional (describing the overall variance explained by the model), and R² marginal (the variance explained only by the fixed effects included in the model), which were obtained with the package *performance* (Lüdecke et al., 2021). Finally, type III ANOVA for linear mixed effect models tests were performed due to the inclusion of the interaction term, to obtain the significance of each predictor included in our best model. Furthermore, using the *emmeans* package (Lenth et al., 2023) we extracted the estimated marginal means (EMMs), confidence intervals (CIs) and significance values of pairwise comparisons (method:tolerance) and we used these data to produce interaction plots in order to observe significant differences in performance for different tolerance categories according to the modelling method used.

## 3. Results

### 3.1. Model results

A total of 62% (n = 523) of the models developed with Period1 (drought-free) met the criteria for being considered as good models (mean AUC ± SD; 0.82 ± 0.08). The evaluation metrics employed to assess the accuracy of model predictions (AUC and TSS) exhibited a strong correlation (Pearson = 0.78, p-value <0.001; Fig. 3a). Wilcoxon tests revealed significant differences between the Period1Acc compared to Period2Acc (p <0.001; Fig. 3). The distribution of the AUCgap showed signs of non-normality (Shapiro test p <0.001). KW tests for Period1Acc showed significant differences according to the method used (p = 0.03) but remained unaffected by the tolerance of species (p = 0.09).

**Fig 3.**
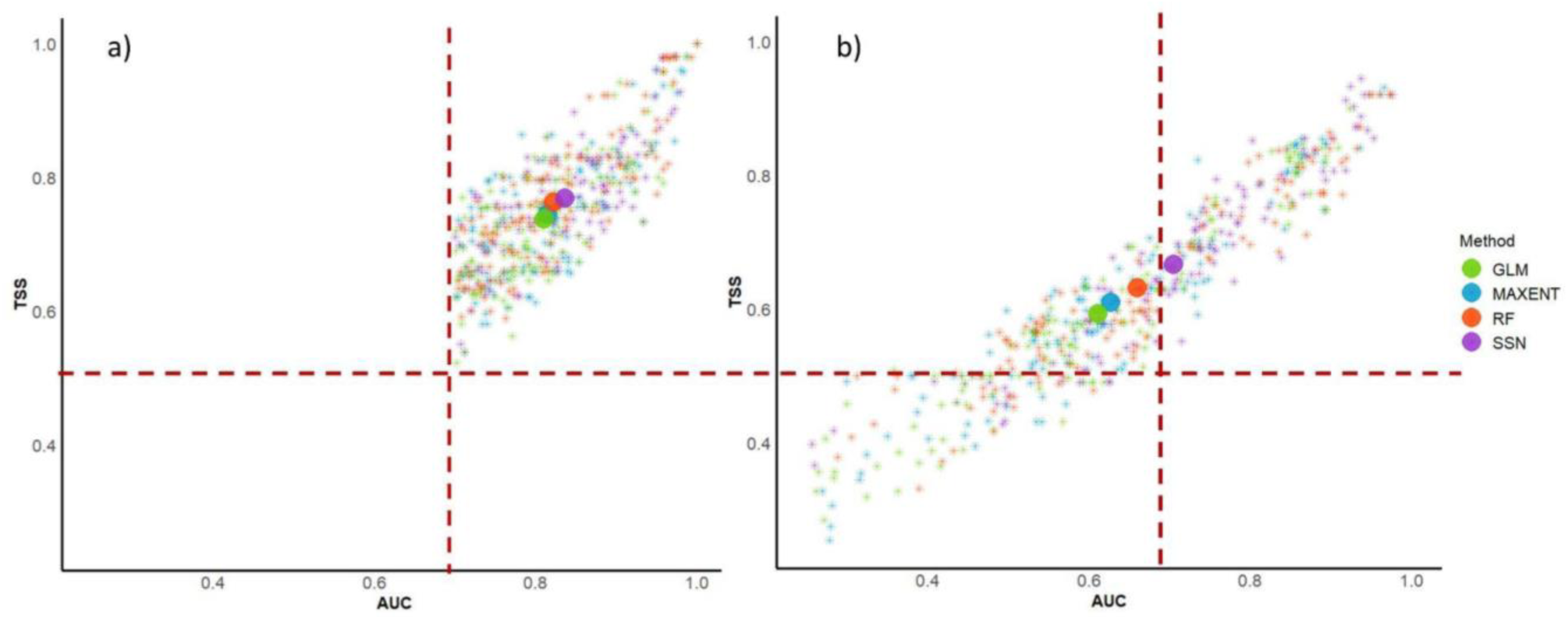
Scores for all 523 valid models in terms of TSS and AUC, for (a) validating species in drought free periods (Period1Acc) and (b) predicting species under the influence of drought (Period2Acc). Dashed lines represent thresholds for fair predictions according to AUC (≥ 0.7) and TSS (≥ 0.5). Stars represent individual models scores and circles represent the mean of all models per modelling method.

From the 523 valid models used to predict species distributions after drought and assessed with presence-absence data from Period2, 39% exhibited good predictive performance (AUC ≥ 0.7; n = 205) and 81% (n = 426) outperformed randomness, successfully predicting over 50% of the data (mean AUC ± SD; 0.65 ± 0.17). AUC scores of predictions under drought influence demonstrated a very strong correlation with TSS (Pearson = 0.94, p <0.001; Fig. 3b) and therefore, further analyses were performed with AUC only. KW test revealed highly significant variations in Period2Acc across the methods used and for different species tolerances (Table S3).

IHAs were more frequently selected than any other variable group for a total of 116 instances (Fig. 4). The most frequently selected single variables were bio1 and elevation chosen on 10 occasions each, followed by Bio19, the upstream percentages of open land and cropland, and ml1 (magnitude of low flows). Variables related to timing and rate of flow events were the least chosen for describing species distributions. In addition, most frequently selected variables showed moderate variability according to drought exposure (Fig. S6)

**Fig. 4.**
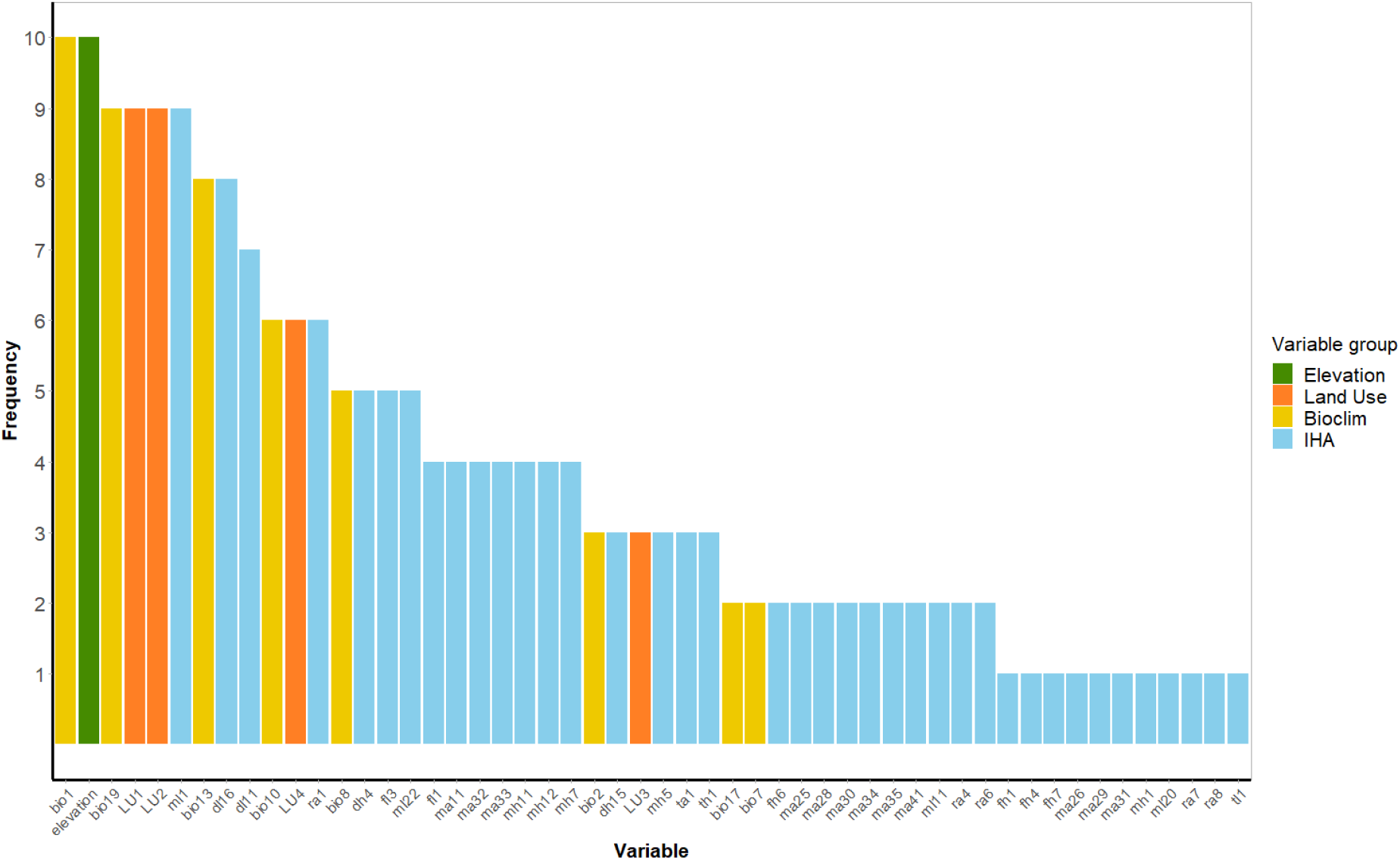
Variable selection frequency. Frequency represented as the number of species including the variable into the model describing their distribution.

### 3.2. Accuracy gap analysis

The predictive accuracy of models deemed "good" on Period1 displayed moderate variation when used to predict the presence and absences of macroinvertebrates during Period2, with a mean accuracy deviation of around 21% in AUC (mean AUCgap ± SD; 0.17 ± 0.17). Simple linear models revealed a significant trend between Period1Acc and the AUCgap they obtained (R² = 0.051, p <0.001; Fig. S4), as well as a significant relationship between the AUCgap and the prevalence of the species in the dataset (R² = 0.013, p = 0.005; Fig. S5). The mean number of occurrences (+SD) of tolerant (47.32 ± 29.02) and sensitive (40.13 ± 19.40) species, was higher than for moderately tolerant (32.52 ± 5.62) and moderately sensitive (35.71 ± 26.17) species, although prevalence was variable during years (Fig. S7).

We observed a strong influence of the tolerance score of the species modelled in the AUCgap between Period1Acc and Period2Acc (KW p <0.001; Fig. 5). Post-hoc Dunn tests showed that tolerant species (0.23 ± 0.17 SD) encountered statistically bigger AUC differences when compared to moderately tolerant (0.09 ± 0.17, p <0.001), moderately sensitive species (0.08 ± 0.1, p <0.001), and when compared to sensitive species (0.16 ± 0.13, p = 0.005). Sensitive species showed higher AUCgap than moderately sensitive (p <0.001) and moderately tolerant species (p = 0.003). The difference in AUCgap between moderately sensitive and moderately tolerant species (p = 0.27) was not significant. Additionally, KW along with Dunn post-hoc test revealed moderate variability in AUCgap based on the method used (p = 0.015; Fig. 6). SSNs (mean AUCgap ± SD; 0.13 ± 0.15) showed lower AUCgap compared to GLMs (0.20 ± 0.18, p = 0.007) and MaxEnt (0.19 ± 0.18, p = 0.017) but not when compared to RFs (0.16 ± 0.17, p= 0.068), no other significant differences between groups were found (p >0.05).

**Fig 5.**
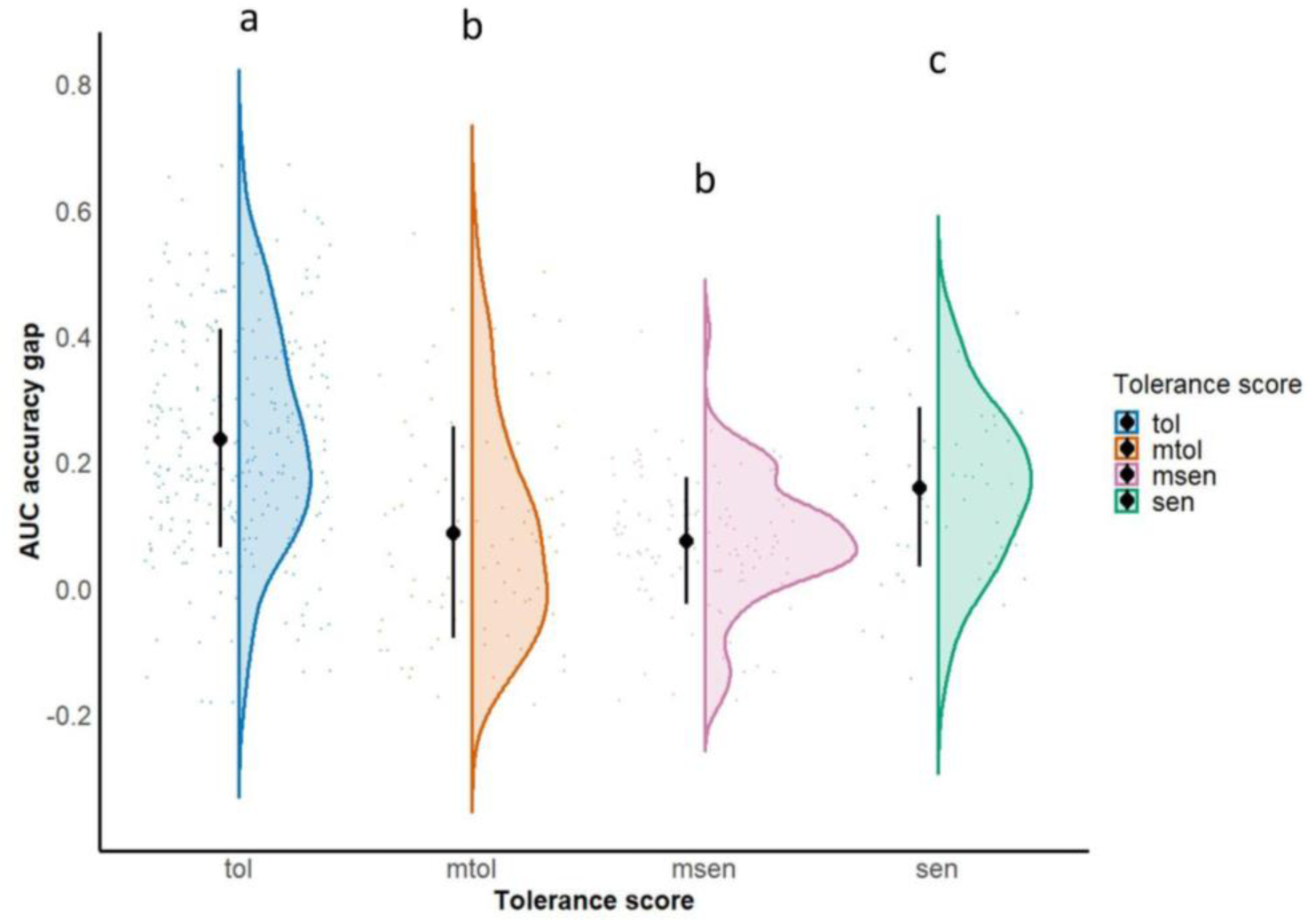
AUCgap (y-axis), according to the tolerance score of the species (x-axis) arranged from more tolerant to more sensitive, categories which do not share letters represent significant differences (alpha <0.05) between them according to Dunn post-hoc tests. Black dots represent means and error bars represent SD; tol = tolerant (n = 23 species; 273 models), mtol = moderately tolerant (n = 6 species; 88 models), msen = moderately sensitive (n = 7 species; 109 models), sen = sensitive (n = 4 species, 53 models).

**Fig 6.**
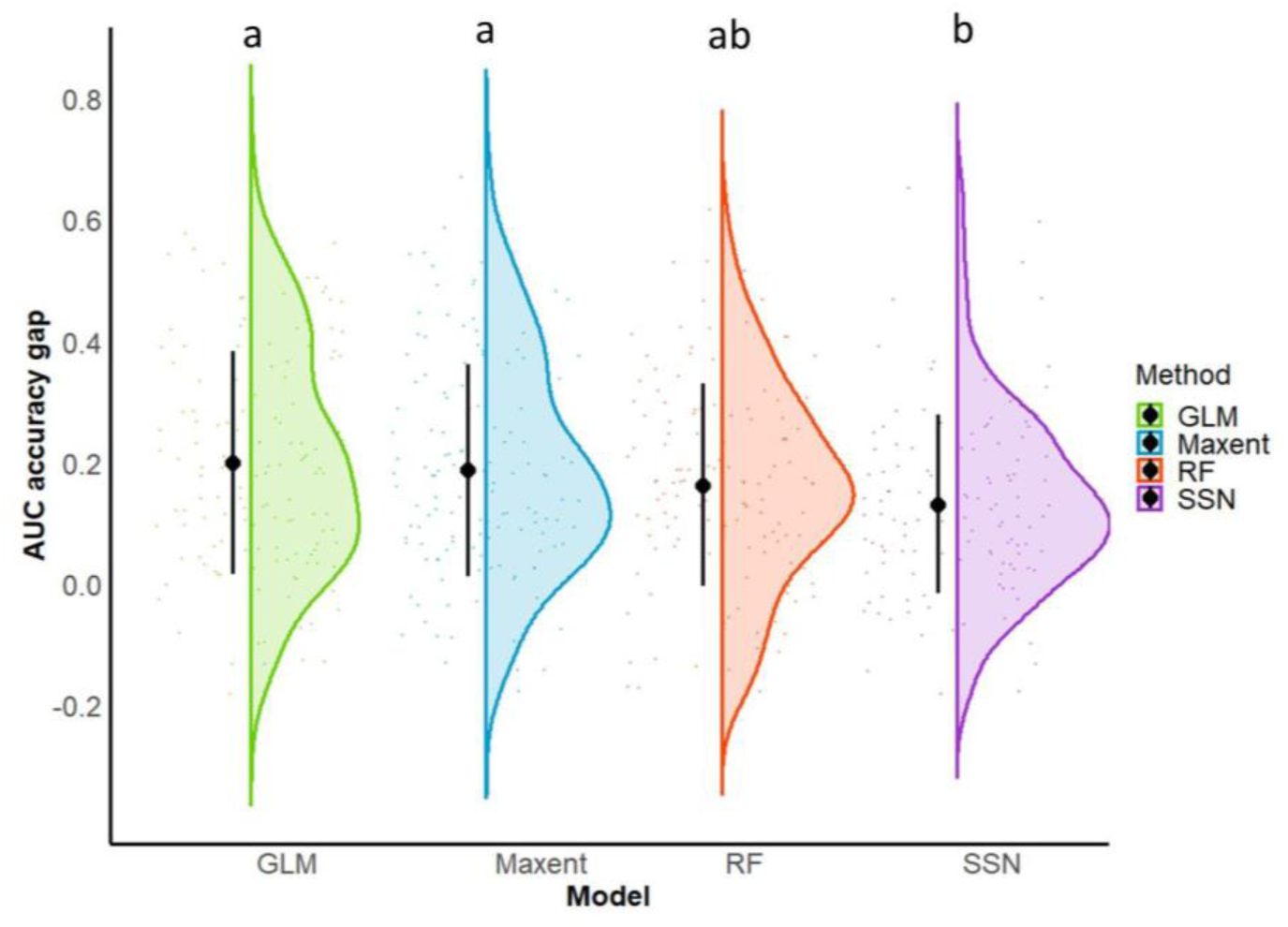
AUCgap (y-axis) depending on the modelling method used (x-axis), categories which do not share letters represent significant differences (alpha <0.05) between them according to Dunn post-hoc tests. Black dots represent means and error bars represent SD.

High variability in AUCgap depending on the species modelled was visually observed (Fig. S3) and was confirmed with a random effects model (Formula 1) with a high variance explained uniquely with this term (R² = 0.72). Moreover, additional variance was explained by the inclusion of tolerance score, modelling method and the interaction between them as fixed effects. Our full model (Formula 2) obtained the best performance in terms of AIC and R² (Table 2).

**Table 2.** Ranking of the models fitted to describe the differences in AUCgap. Ranking represents the position of the model from best to worst performance, and was obtained by comparing several metrics. R² conditional describes the overall variance explained by the model, R² marginal is the variance explained only by the fixed effects included in the model. AICc weights describe the relative likelihood of models being the most parsimonious.

| Ranking | Model | R<sup>2</sup> conditional | R<sup>2</sup> marginal | AICc weights |
| --- | --- | --- | --- | --- |
| 1 | Tolerance*Method | 0.760 | 0.165 | 0.518 |
| 2 | Tolerance+Method | 0.752 | 0.155 | 0.356 |
| 3 | Tolerance | 0.738 | 0.141 | <0.001 |
| 4 | Method | 0.739 | 0.017 | 0.126 |
| 5 | Random model only | 0.725 | 0.000 | <0.001 |

The type III ANOVA tests for the full model (Table S4) highlighted the significance of the modelling method (p <0.001) and tolerance scores (p = 0.040) to determine the accuracy gap between models before and after the occurrence of drought. Moreover, interaction derived through type III ANOVA was also significant (p = 0.028). In addition, estimated marginal means revealed that sensitive species models performed notably better when using SSNs, showing significantly lower accuracy gaps than when modelled with GLMs MaxEnt and RFs (Fig. 7). Tolerant species demonstrated lower AUCgap when modelled using SSNs and RFs. Conversely, pairwise comparisons across modelling methods and for moderately tolerant and moderately sensitive species did not yield significant differences, suggesting no substantial reduction in errors for these species, regardless of the model used.

**Fig 7.**
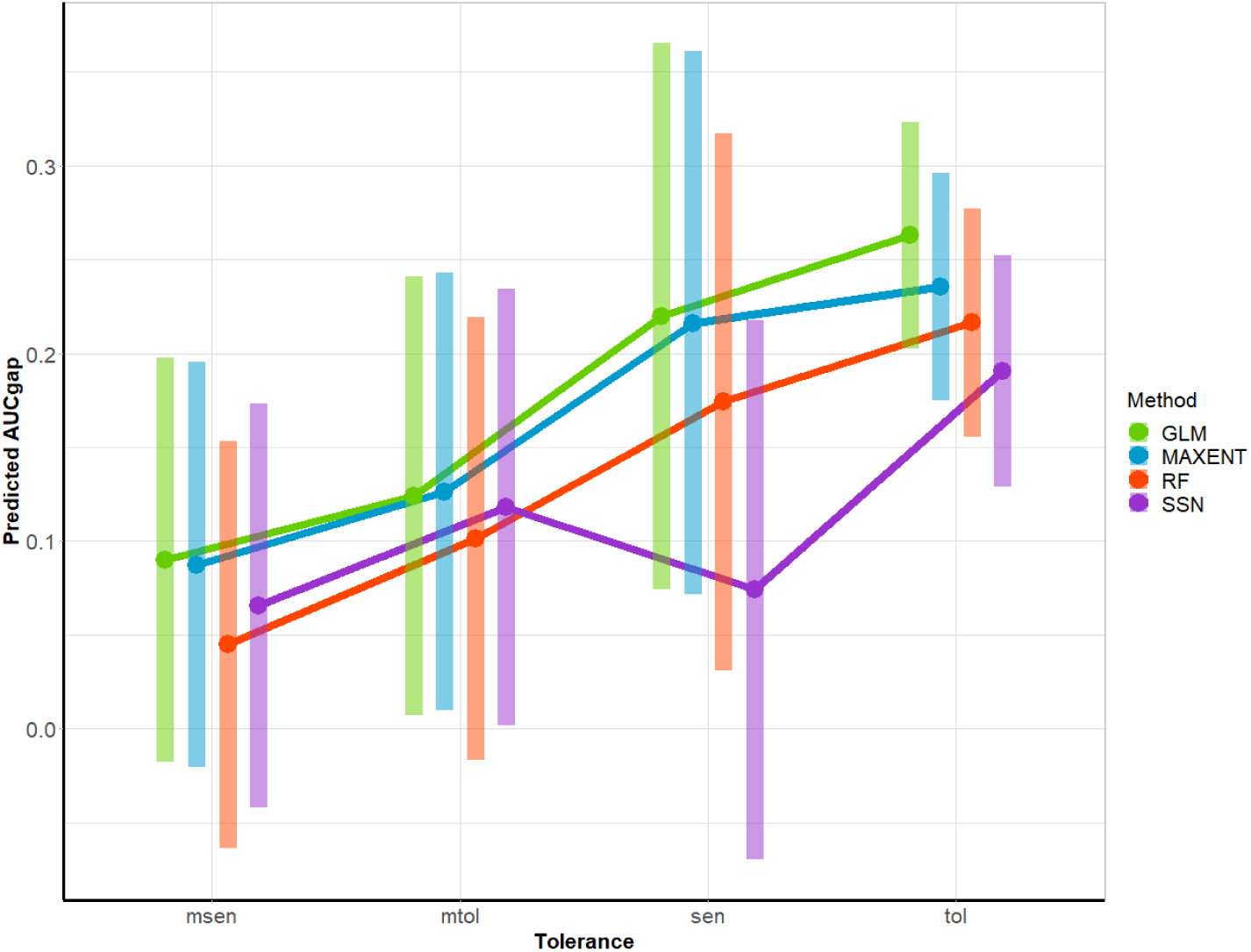
Interaction plot illustrating estimated marginal means derived from the full model. The y-axis depicts the predicted values of AUCgap, categorised by both the tolerance score of the species (on the x-axis) and the modelling method employed. The X-axis is arranged in ascending order of mean AUCgap values and shaded areas represent 95% CIs.

## 4. Discussion

### 4.1 Are SDMs from drought-free conditions transferable to drought-influenced conditions?

Our study revealed noticeable differences in the predictive accuracy of models differing in drought exposure. While previous research assessing SDM transferability to drought conditions is lacking, studies across various taxonomic groups (Nguyen et al., 2022; Gies et al., 2015) have shown reduced accuracies when transferred to novel environments, indicating that the accuracy of SDMS obtained through internal validation, may not indicate their transferability to different contexts (Petitpierre et al., 2017). Notably, our findings highlight that SDM transferability might fluctuate according to the tolerance score of the modelled species and the modelling method used.

The limited transferability of SDMS from drought-free conditions to drought-influenced ones might stem from environmental novelty associated with drought (Feng et al., 2019; Gies et al., 2015; Owens et al., 2013), and the models’ inability to generalise responses beyond training data. Extrapolating beyond training conditions is necessary when predicting species’ distributions under extreme event conditions (Franklin, 2023), hence, understanding SDM transferability becomes crucial. Restricted SDM transferability may have been a result of shifting variable relevance for the species under drought exposure (Feng et al., 2019). Without drought, certain environmental variables may be identified as influential factors for species distributions. However, during droughts, the influence of these variables can change, potentially reducing the accuracy of the SDMs (Rousseau & Betts, 2022; Owens et al., 2013). In our study, frequently selected variables (e.g. precipitation of the coldest and wettest months, mean minimum monthly flows and mean temperatures of the warmest quarter and wettest quarter) showed different ranges under drought exposure, possibly reducing SDM transferability. Additionally, during droughts, besides abiotic factors, biotic interactions may increasingly influence the distribution of freshwater macroinvertebrates (Bond et al., 2008; Dormann et al., 2018). For instance, droughts might translate to a reduced flow velocity and habitat availability, increasing the competition and predator-prey interactions (Dewson, James & Death, 2007). Moreover, after flow resumption, dispersal occurs (Tonkin et al., 2018) and priority effects can define the structure of the community by shaping biotic interactions (Little & Altermatt, 2018). Therefore, incorporating these mechanisms might serve to enhance the reliability of predictive models.

### 4.2. Do SDMs based on specific tolerance scores yield higher transferability?

Our findings suggest that models used for predicting either tolerant or sensitive species during exposure to droughts might show lower transferability in comparison to moderately tolerant and moderately sensitive species. Tolerant species may exhibit lower transferability, as indicated by studies highlighting the dependency of SDM accuracy on species’ ecology, particularly regarding species with broader ranges (Wogan, 2016; Grenouillet et al., 2011; Rousseau & Betts, 2022) and the inability of models to effectively determine their environmental niche. Nevertheless, sensitive species also showed high accuracy gaps, contrary to the suggestion that species with narrower ranges might be easier to predict, this result might be related to the low number of species with a sensitive score (n = 4). In addition, the higher prevalence of sensitive and tolerant species could have affected the ability of the model to discriminate from suitable and unsuitable habitats.

Moreover, droughts can influence abiotic processes, resulting in changes in water quality such as elevated salinity, decreased dissolved oxygen levels, and shifts in nutrient availability in rivers (Mosley, 2015), exceeding critical thresholds within ecological communities (Bailey & van de Pol, 2016). In this context, tolerant species may show nonspecific responses to stressors, due to larger withstand thresholds, which SDMs might not fully capture (Thuiller et al., 2004; Guevara et al., 2018). Although the prevalence of sensitive species was relatively lower than for tolerant species, it showed more fluctuations, probably associated with environmental changes related to droughts, possibly affecting model accuracy. In fact, the combined impact of stressors on freshwater macroinvertebrate distribution has been acknowledged (Mantyka-Pringle et al., 2014), but further exploration is warranted to assess how drought interacts with other environmental variables, possibly affecting the accuracy of SDMs.

Species in the moderate range of the tolerance spectrum (moderately tolerant and moderately sensitive) showed higher transferability. This might have been due to their median tolerances resulting in less exaggerated responses to drought, thereby adhering more closely to the distribution outlined by the initial SDMs. In addition, the prevalence of these species in our dataset was lower, which could have helped to define their environmental preferences (Grenouillet et al., 2011). Our findings underscore the pivotal role of species tolerance in determining their distribution post-drought, highlighting its potential contribution to shaping communities amidst heightened occurrences of stress pulses. Nonetheless, we acknowledge that the tolerance score utilised in this study is limited to stream zonation traits. Thus, although using the tolerance score remains a practical tool for biomonitoring, the use of other influential traits affecting species distribution under hydrological stress (e.g. current preference and dispersal modes) warrants further investigation.

### 4.3. Are there modelling methods that are more transferable?

Our findings revealed that SDM transferability might be impacted by the modelling method used, agreeing with other studies examining SDM spatial transferability (Charney et al., 2021). In particular, SSNs showed higher transferability to conditions under drought exposure, when compared with GLM and MaxEnt models. Our research highlights the potential value of integrating SSNs into predictive modelling within freshwater ecosystems, particularly in ecological settings where connectivity is crucial for predicting species occurrence. Lower transferability obtained by presence only approaches (MaxEnt) and GLMs, aligns with prior research indicating that MaxEnt can be negatively affected by environmental novelty (Feng et al., 2019). The similar transferability obtained by MaxEnt, RF and GLM models resonates with broader trends observed in analogous studies, where models of species present in more than 25% of the dataset reported little to no difference in accuracy according to the method used (Benkendorf et al., 2023). In our study we observed a slight but significant effect of prevalence on transferability, similar to previous studies (Gies et al., 2015). Therefore, we suggest that further studies focus on exploring whether class imbalances (e.g. presence/absence) might have an impact on the transferability of models to novel conditions.

### 4.4. Does any modelling method transfer better to drought-influenced conditions for certain tolerance scores?

Our study showed that the transferability of models for sensitive species was enhanced by using SSNs, and for tolerant species using SSNs and RFs. These results highlight the need to incorporate models that consider the ecological complexities of freshwater systems under hydrological stress. For instance, RFs, in comparison with other methods, excel in handling complex variable interactions (Merow et al., 2014; Elith et al., 2006) and are able to better capture stressor interactions affecting sensitive species. While the veracity of this remains uncertain, investigating methods proficient in capturing variable interactions (Golding et al., 2016) and delving into stressor interaction mechanisms could enhance species distribution predictions, especially in extreme flow contexts.

In addition, aside from the environmental predictors commonly included in SDMs, additional processes such as dispersal, can shape species distributions (Sarremejane et al., 2017). These processes may exhibit spatially autocorrelated patterns that are not adequately captured by standard SDMs (Merow et al., 2014; Barry & Elith, 2006) and which could become more significant in the context of habitat fragmentation resulting from droughts (Robson et al., 2011; Aspin et al., 2019a; Aspin et al., 2019b). In such cases, connectivity becomes essential as species might rely on finding refugia to survive droughts or repopulate areas after such events (Robson et al., 2011). SSNs may be particularly effective in predicting sensitive species with limited distributions by confining predictions to areas with connections to observed occurrences, thus enhancing accuracy. Although our sample size for sensitive species models is relatively small (n = 53) compared to other tolerance scores, our results underscore the importance of accounting for stream connectivity in the estimation of the impacts of climate change. Particularly for sensitive species, access to refugia for species unable to disperse rapidly under these conditions could be crucial in determining their persistence or local extinction from the system.

## 5. Conclusion

SDMs have evolved into indispensable tools for ecological research, especially in assessing how climate change might impact species distribution. Nonetheless, the application of SDMs in predicting species responses during extreme flows, like droughts, has received limited attention. This knowledge gap is particularly critical in freshwater ecosystems, where droughts are known to pose significant threats to riverine species. Our study highlights that the transferability of SDMs to drought conditions can be affected by the tolerance of the species and the chosen modelling method, as well as the relevance of particular modelling methods for modelling specific species. Recognizing the non-stationarity in species responses, and accounting for the connectivity of river ecosystems are essential for accurate predictions in the face of climate change-induced stress. Furthermore, as the quantity and diversity of stressors in freshwater ecosystems continue to increase, we suggest that future studies on SDM transferability should focus on understanding the impact of extreme flows (and other extreme events), coupled with ongoing environmental alterations related to climate change, and the interplay of biotic processes on our capacity to predict species distributions accurately across varied contexts. Incorporating these nuances into SDMs is crucial for better anticipating and managing the impacts of stress pulses on riverine species, such as those produced by droughts.

## Notes

### Competing Interest Statement

The authors have declared no competing interest.

## References

Allouche, O., Tsoar, A., & Kadmon, R. (2006). Assessing the accuracy of species distribution models: prevalence, kappa and the true skill statistic (TSS). Journal of applied ecology, 43(6), 1223–1232.

Altermatt, F. (2013). Diversity in riverine metacommunities: a network perspective. Aquatic Ecology, 47(1), 365–377. 10.1007/s10452-013-9450-3

Aspin, T.W.H., Hart, K., Khamis, K., et al. (2019). Drought intensification alters the composition, body size, and trophic structure of invertebrate assemblages in a stream mesocosm experiment. Freshwater Biology; 64: 750–760. 10.1111/fwb.13259

Aspin, T.W.H., Khamis, K., Matthews, T.J., et al. (2019). Extreme drought pushes stream invertebrate communities over functional thresholds. Global Change Biology, 25(1), 230–244. 10.1111/gcb.14495

Bailey, L. D., & van de Pol, M. (2016). Tackling extremes: challenges for ecological and evolutionary research on extreme climatic events. Journal of Animal Ecology, 85(1), 85–96.

Baldan, D., Kiesel, J., Hauer, C., Jähnig, S.C., Hein, T. (2021). Increased sediment deposition triggered by climate change impacts Freshwater Pearl Mussels habitats and metapopulations. Journal of Applied Ecology, 58 (9),1933–1944. 10.1111/1365-2664.13940

Barry, S., & Elith, J. (2006). Error and uncertainty in habitat models. Journal of Applied Ecology, 43(3), 413–423.

Bates, D., Mächler, M., Bolker, B., & Walker, S. (2015). Fitting linear mixed-effects models using lme4. Journal of Statistical Software, 67(1), 1–48. 10.18637/jss.v067.i01

Beniston, M., Stephenson, D.B., Christensen, O.B. et al. (2007). Future extreme events in European climate: an exploration of regional climate model projections. Climatic Change 81(1), 71–95. 10.1007/s10584-006-9226-z

Benkendorf, D. J., Schwartz, S. D., Cutler, D. R., & Hawkins, C. P. (2023). Correcting for the effects of class imbalance improves the performance of machine-learning based species distribution models. Ecological Modelling, 483, 110414. 10.1016/j.ecolmodel.2023.110414.

Bertoncin, A.P.D.S., Pinha, G.D., Baumgartner, M.T., & Mormul, R.P. (2019). Extreme drought events can promote homogenization of benthic macroinvertebrate assemblages in a floodplain pond in Brazil. Hydrobiologia, 826(1), 379–393. 10.1007/s10750-018-3756-z

Bond, N. R., Lake, P. S., & Arthington, A. H. (2008). The impacts of drought on freshwater ecosystems: an Australian perspective. Hydrobiologia, 600(1), 3–16. 10.1007/s10750-008-9326-z

Boulton, A.J. (2003), Parallels and contrasts in the effects of drought on stream macroinvertebrate assemblages. Freshwater Biology, 48(1), 1173–1185. 10.1046/j.1365-2427.2003.01084.x

Calapez, A. R., Elias, C. L., Almeida, S. F., & Feio, M. J. (2014). Extreme drought effects and recovery patterns in the benthic communities of temperate streams. Limnetica, 33(2), 281–296. 10.23818/limn.33.22

Charney, N. D., Record, S., Gerstner, B. E., Merow, C., Zarnetske, P. L., & Enquist, B. J. (2021). A test of species distribution model transferability across environmental and geographic space for 108 western North American tree species. Frontiers in Ecology and Evolution, 9, 689295. 10.3389/fevo.2021.689295

Chessman, B.C. (2015). Relationships between lotic macroinvertebrate traits and responses to extreme drought. Freshwater Biology, 60(1), 50–63. 10.1111/fwb.12466

Collart, F., Broennimann, O., Guisan, A. and Vanderpoorten, A. (2023). Ecological and biological indicators of the accuracy of species distribution models: lessons from European bryophytes. Ecography, 2023(8), e06721. 10.1111/ecog.06721

Comte, L., Olden, J. (2017). Climatic vulnerability of the world’s freshwater and marine fishes. Nature Climate Change, 7(1), 718–722. 10.1038/nclimate3382

Daufresne, M., Bady, P., & Fruget, J. F. (2007). Impacts of global changes and extreme hydroclimatic events on macroinvertebrate community structures in the French Rhône River. Oecologia, 151(1), 544–559. 10.1007/s00442-006-0655-1

Dewson, Z. S., James, A. B., & Death, R. G. (2007). A review of the consequences of decreased flow for instream habitat and macroinvertebrates. Journal of the North American Benthological Society, 26(3), 401–415. 10.1899/06-110.1

Domisch, S., Araújo, M.B., Bonada, N., Pauls, S.U., Jähnig, S.C. and Haase, P. (2013). Modelling distribution in European stream macroinvertebrates under future climates. Global Change Biology, 19(3), 752–762. 10.1111/gcb.12107

Dormann, C. F. et al. (2018). Biotic interactions in species distribution modelling: 10 questions to guide interpretation and avoid false conclusions. Global Ecology and Biogeography, 27(9), 1004– 1016. 10.1111/geb.12759.

Eger, A. M., Curtis, J. M., Fortin, M. J., Côté, I. M., & Guichard, F. (2017). Transferability and scalability of species distribution models: a test with sedentary marine invertebrates. Canadian Journal of Fisheries and Aquatic Sciences, 74(5), 766–778. 10.1139/cjfas-2016-0129

Elith, J., et al. (2006). Novel methods improve prediction of species’ distributions from occurrence data. Ecography, 29(2), 129–151. 10.1111/j.2006.0906-7590.04596.x

Elo, M., Jyrkänkallio-Mikkola, J., Ovaskainen, O., Soininen, J., Tolonen, K.T., Heino, J. (2021), Does trait-based joint species distribution modelling reveal the signature of competition in stream macroinvertebrate communities? Journal of Animal Ecology, 90(5), 1276–1287. 10.1111/1365-2656.13453

Feng, X., Park, D. S., Liang, Y., Pandey, R., & Papeş, M. (2019). Collinearity in ecological niche modeling: Confusions and challenges. Ecology and evolution, 9(18), 10365–10376. 10.1002/ece3.5555

Fortuna, M. A., Gómez-Rodríguez, C., & Bascompte, J. (2006). Spatial network structure and amphibian persistence in stochastic environments. Proceedings of the Royal Society B: Biological Sciences, 273(1592), 1429–1434. 10.1098/rspb.2005.3448

Franklin, J. (2023). Species distribution modelling supports the study of past, present and future biogeographies. Journal of Biogeography, 50(9), 1533–1545. 10.1111/jbi.14617

Friedman, J., Hastie, T., & Tibshirani, R. (2010). Regularization paths for generalized linear models via coordinate descent. Journal of statistical software, 33(1), 1–22. 10.18637/jss.v033.i01

Gies, M., Sondermann, M., Hering, D., & Feld, C. K. (2015). Are species distribution models based on broad-scale environmental variables transferable across adjacent watersheds? A case study with eleven macroinvertebrate species. Fundamental and Applied Limnology, 186(1-2), 63–97. 10.1127/fal/2014/0600

Golding, N., & Purse, B. V. (2016). Fast and flexible Bayesian species distribution modelling using Gaussian processes. Methods in Ecology and Evolution, 7(5), 598–608. 10.1111/2041-210X.12523

Graf, W., Zoltan, L., Leitner, P., Dossi, F. (2018). Simplifying biotic assessment – a novel Tolerance Index for macro-invertebrate communities. University for Natural Resources and Life Sciences, Vienna, Department of Water, Atmosphere and Environment Institute of Hydrobiology and Aquatic Ecosystem Management.

Grenouillet, G., Buisson, L., Casajus, N., & Lek, S. (2011). Ensemble modelling of species distribution: the effects of geographical and environmental ranges. Ecography, 34(1), 9–17. 10.1111/j.1600-0587.2010.06152.x

Guevara, L., Gerstner, B. E., Kass, J. M., & Anderson, R. P. (2018). Toward ecologically realistic predictions of species distributions: A cross-time example from tropical montane cloud forests. Global Change Biology, 24(4), 1511–1522.. 10.1111/gcb.13992.

Guisan, A., Thuiller, W., & Zimmermann, N. E. (2017). Habitat suitability and distribution models: with applications in R. Cambridge University Press.

Guisan, A. et al. (2013). Predicting species distributions for conservation decisions. Ecology letters, 16(12), 1424–1435. 10.1111/ele.12189

Haase, P., Pilotto, F., Li, F., Sundermann, A., Lorenz, A. W., Tonkin, J. D., & Stoll, S. (2019). Moderate warming over the past 25 years has already reorganized stream invertebrate communities. Science of the Total Environment, 658, 1531–1538. 10.1016/j.scitotenv.2018.12.234

Haase, P., Sundermann, A., Schindehütte, K. (2006) Informationstext zur Operationellen Taxaliste als Mindestanforderung an die Bestimmung von Makrozoobenthosproben aus Fließgewässern zur Umsetzung der EU-Wasserrahmenrichtlinie in Deutschland. https://gewaesser-bewertung-berechnung.de/files/downloads/perlodes/Operationelle_Taxaliste_Begleittext.pdf.

He, F., et al. (2019). The global decline of freshwater megafauna. Global Change Biology, 25(11), 3883–3892. 10.1111/gcb.14753

Herbst, D. B., Cooper, S. D., Medhurst, R. B., Wiseman, S. W., & Hunsaker, C. T. (2019). Drought ecohydrology alters the structure and function of benthic invertebrate communities in mountain streams. Freshwater Biology, 64(5), 886–902. 10.1111/fwb.13270

Hernandez, P.A., Graham, C.H., Master, L.L. and Albert, D.L. (2006), The effect of sample size and species characteristics on performance of different species distribution modeling methods. Ecography, 29(5), 773–785. 10.1111/j.0906-7590.2006.04700.x

Hill, M.J., Mathers, K.L., Little, S. et al. (2019), Ecological effects of a supra-seasonal drought on macroinvertebrate communities differ between near-perennial and ephemeral river reaches. Aquatic Sciences 81, 62. 10.1007/s00027-019-0659-7

IPCC. (2023). AR6 Synthesis Report. Climate change. https://www.ipcc.ch/report/ar6/syr/. Accessed on March 31st 2024.

Irving, K., Jähnig, S. C., & Kuemmerlen, M. (2022). Disentangling the effect of climatic and hydrological predictor variables on benthic macroinvertebrate distributions from predictive models. Hydrobiologia, 849(4), 1021–1040. 10.1007/s10750-021-04765-w

Kakouei K., Kiesel J., Domisch S., Irving K.S., Jähnig S.C., Kail J. (2018). Projected effects of climate change-induced flow alterations on stream macroinvertebrate abundances. Ecology and Evolution, 8(6), 3393–3409. 10.1002/ece3.3907

King, A. J., Tonkin, Z., & Lieshcke, J. (2012). Short-term effects of a prolonged blackwater event on aquatic fauna in the Murray River, Australia: considerations for future events. Marine and Freshwater Research, 63(7), 576–586. 10.1071/MF11275

Konowalik, K., & Nosol, A. (2021). Evaluation metrics and validation of presence-only species distribution models based on distributional maps with varying coverage. Scientific Reports, 11(1), 1482. 10.1038/s41598-020-80062-1

Kuemmerlen, M., Schmalz, B., Guse, B., Cai, Q., Fohrer, N., & Jähnig, S. C. (2014). Integrating catchment properties in small scale species distribution models of stream macroinvertebrates. Ecological Modelling, 277, 77–86. 10.1016/j.ecolmodel.2014.01.020

Kundzewicz, Z. et al. (2014). Flood risk and climate change: global and regional perspectives. Hydrological Sciences Journal, 59(1), 1–28. 10.1080/02626667.2013.857411

Lenth, R. et al. (2023). emmeans: Estimated Marginal Means, aka Least-Squares Means. R package. https://CRAN.R-project.org/package=emmeans

Liaw, A., & Wiener, M. (2002). Classification and regression by randomForest. R news, 2(3), 18–22.

Little, C. J., & Altermatt, F. (2018). Do priority effects outweigh environmental filtering in a guild of dominant freshwater macroinvertebrates?. Proceedings of the Royal Society B: Biological Sciences, 285(1876), 20180205. 10.1098/rspb.2018.0205

Liu, C., Berry, P. M., Dawson, T. P., & Pearson, R. G. (2005). Selecting thresholds of occurrence in the prediction of species distributions. Ecography, 28(3), 385–393. 10.1111/j.0906-7590.2005.03957.x

Lüdecke, D., et al. (2021). performance: An R Package for Assessment, Comparison and Testing of Statistical Models. Journal of Open Source Software, 6(60), 3139. 10.21105/joss.03139

Madsen, P. B., et al. (2015). Altitudinal distribution limits of aquatic macroinvertebrates: an experimental test in a tropical alpine stream. Ecological Entomology, 40(5), 629–638. 10.1111/een.12232

Mantyka-Pringle, C. S., Martin, T. G., Moffatt, D. B., Linke, S., & Rhodes, J. R. (2014). Understanding and predicting the combined effects of climate change and land-use change on freshwater macroinvertebrates and fish. Journal of Applied Ecology, 51(3), 572–581. 10.1111/1365-2664.12236

Markovic, D., Carrizo, S.F., Kärcher, O., Walz, A. and David, J.N.W. (2017), Vulnerability of European freshwater catchments to climate change. Global Change Biology, 23(9), 3567–3580. 10.1111/gcb.13657

McPherson, J. and Jetz, W. (2007), Effects of species’ ecology on the accuracy of distribution models. Ecography, 30(1), 135–151. 10.1111/j.0906-7590.2007.04823.x

Meehl, G. A., & Tebaldi, C. (2004). More intense, more frequent, and longer lasting heat waves in the 21st century. Science, 305(5686), 994–997. 10.1126/science.1098704

Meier, C., Haase, P., Rolauffs, P., Schindehütte, K., Schöll, F., Sundermann, A., & Hering, D. (2006). Methodisches Handbuch Fließgewässerbewertung - Handbuch zur Untersuchung und Bewertung von Fließgewässern auf der Basis des Makrozoobenthos vor dem Hintergrund der EG-Wasserrahmenrichtlinie (Vol. Stand: Mai 2006). https://gewaesser-bewertung-berechnung.de/

Merow, C., et al. (2014). What do we gain from simplicity versus complexity in species distribution models?. Ecography, 37(12), 1267–1281. 10.1111/ecog.00845

Messager, M. L., et al. (2021). Global prevalence of non-perennial rivers and streams. Nature 594, 391–397. 10.1038/s41586-021-03565-5

Miller, J. (2010), Species Distribution Modeling. Geography Compass, 4(6), 490–509. 10.1111/j.1749-8198.2010.00351.x

Moog, O. (1995). Fauna Aquatica Austriaca - A Comprehensive Species Inventory of Austrian Aquatic Organisms with Ecological Notes. Federal Ministry for Agriculture and Forestry, Wasserwirtschaftskataster Vienna.

Mosley, L. M. (2015). Drought impacts on the water quality of freshwater systems; review and integration. Earth-Science Reviews, 140(1), 203–214. 10.1016/j.earscirev.2014.11.010

Nguyen, D., & Leung, B. (2022). How well do species distribution models predict occurrences in exotic ranges?. Global Ecology and Biogeography, 31(6), 1051–1065. 10.1111/geb.13482

Nguyen, H.H., Kiesel, J., Peters, K. et al. (2023). Stream macroinvertebrate community metrics consistently respond to a spatiotemporal disturbance gradient but composition is more context-dependent. Landscape Ecology, 38 (1), 3133–3151. 10.1007/s10980-023-01769-w

Nguyen, H. H., et al. (2024). Stream macroinvertebrate communities in restored and impacted catchments respond differently to climate, land-use, and runoff over a decade. Science of The Total Environment, 929, 172659. 10.1016/j.scitotenv.2024.172659

Nilsson, C., Polvi, L.E. and Lind, L. (2015), Extreme events in streams and rivers in arctic and subarctic regions in an uncertain future. Freshwater Biology, 60(12), 2535–2546. 10.1111/fwb.12477

Olden, J.D. & Poff, N.L. (2003). Redundancy and the choice of hydrologic indices for characterizing streamflow regimes. River Research and Applications, 19(2), 101–121. 10.1002/rra.700

Owens, H. L., et al. (2013). Constraints on interpretation of ecological niche models by limited environmental ranges on calibration areas. Ecological modelling, 263, 10–18. 10.1016/j.ecolmodel.2013.04.011

Peterson, A. T., Papeş, M., & Soberón, J. (2015). Mechanistic and Correlative Models of Ecological Niches. European Journal of Ecology, 1(2), 28–38. 10.1515/eje-2015-0014

Petitpierre, B., Broennimann, O., Kueffer, C., Daehler, C., & Guisan, A. (2017). Selecting predictors to maximize the transferability of species distribution models: Lessons from cross-continental plant invasions. Global Ecology and Biogeography, 26(3), 275–287. 10.1111/geb.12530

Phillips, S. J., Dudík, M., & Schapire, R. E. (2004). A maximum entropy approach to species distribution modeling. In Proceedings of the twenty-first international conference on Machine learning, 665–662.

Qiao, H., Feng, X., Escobar, L.E., Peterson, A.T., Soberón, J., Zhu, G. & Papeş, M. (2019). An evaluation of transferability of ecological niche models. Ecography 42(3), 521–534. 10.1111/ecog.03986

R Core Team (2023). R: A language and environment for statistical computing. R Foundation for Statistical Computing, Vienna, Austria. https://www.R-project.org/

Richter, B. D., Baumgartner, J. V., Powell, J., & Braun, D. P. (1996). A method for assessing hydrologic alteration within ecosystems. Conservation biology, 10(4), 1163–1174. 10.1046/j.1523-1739.1996.10041163.x

Robson, B. J., Chester, E. T., & Austin, C. M. (2011). Why life history information matters: drought refuges and macroinvertebrate persistence in non-perennial streams subject to a drier climate. Marine and Freshwater Research, 62(7), 801–810. 10.1071/MF10062

Röpke, C. P., et al. (2017). Simultaneous abrupt shifts in hydrology and fish assemblage structure in a floodplain lake in the central Amazon. Scientific Reports, 7(1), 40170. 10.1038/srep40170

Rousseau, J.S., Betts, M.G. (2022), Factors influencing transferability in species distribution models. Ecography, 2022(7), e06060. 10.1111/ecog.06060

Rückauer, B., Känzig, N., Liu, S. C., Delbruck, T., & Sandamirskaya, Y. (2019). Closing the accuracy gap in an event-based visual recognition task. 10.48550/arXiv.1906.08859

Ruiz, T., et al. (2022). Asynchronous recovery of predators and prey conditions resilience to drought in a neotropical ecosystem. Scientific Reports 12, 8392. 10.1038/s41598-022-12537-2

Sabater, S., et al. (2023). Extreme weather events threaten biodiversity and functions of river ecosystems: evidence from a meta-analysis. Biological Reviews, 98(2), 450–461. 10.1111/brv.12914

Sarremejane, R., Mykrä, H., Bonada, N., Aroviita, J., Muotka. T. (2017). Habitat connectivity and dispersal ability drive the assembly mechanisms of macroinvertebrate communities in river networks. Freshwater Biology, 62(6), 1073–1082. 10.1111/fwb.12926

Schmidt-Kloiber, A. & Hering D. (2015). www.freshwaterecology.info - an online tool that unifies, standardises and codifies more than 20,000 European freshwater organisms and their ecological preferences. Ecological Indicators. 10.1016/j.ecolind.2015.02.007

Schürings, C., Feld, C. K., Kail, J., & Hering, D. (2022). Effects of agricultural land use on river biota: a meta-analysis. Environmental Sciences Europe, 34(1), 124. 10.1186/s12302-022-00706-z

Shah, D.N., Domisch, S., Pauls, S.U., Haase, P., & Jähnig, S.C. (2014). Current and future latitudinal gradients in stream macroinvertebrate richness across North America. Freshwater Science, 33(4), 1136–1147. 10.1086/678492

Shapiro, S. S., & Wilk, M. B. (1965). An analysis of variance test for normality (complete samples). Biometrika, 52(3–4), 591–611. 10.1093/biomet/52.3-4.591

Tessarolo, G., Lobo, J. M., Rangel, T. F., & Hortal, J. (2021). High uncertainty in the effects of data characteristics on the performance of species distribution models. Ecological Indicators, 121, 107147. 10.1016/j.ecolind.2020.107147

Thuiller, W., Brotons, L., Araújo, M. B., & Lavorel, S. (2004). Effects of restricting environmental range of data to project current and future species distributions. Ecography, 27(2), 165–172. 10.1111/j.0906-7590.2004.03673.x

Tickner, D., et al. (2020). Bending the curve of global freshwater biodiversity loss: An emergency recovery plan. BioScience, 70(4), 330–342. 10.1093/biosci/biaa002

Tonkin J.D., Altermatt F, Finn D., et al. (2018), The role of dispersal in river network metacommunities: Patterns, processes, and pathways. Freshwater Biology, 63(1), 141–163. 10.1111/fwb.13037

Van Looy K., et al. (2019) The three Rs of river ecosystem resilience: Resources, Recruitment and Refugia. River Research and Applications, 35(2),107–120. 10.1002/rra.3396

Ver Hoef, J. M., Peterson, E., & Theobald, D. (2006). Spatial statistical models that use flow and stream distance. Environmental and Ecological statistics, 13, 449–464. version 1.8.9. 10.1007/s10651-006-0022-8

Werkowska, W., Márquez, A. L., Real, R., & Acevedo, P. (2017). A practical overview of transferability in species distribution modeling. Environmental reviews, 25(1), 127–133. 10.1139/er-2016-0045

Wogan, G.O. (2016). Life history traits and niche instability impact accuracy and temporal transferability for historically calibrated distribution models of North American birds. PLoS One, 11(3), e0151024. 10.1371/journal.pone.0151024

WWF. (2022), Living Planet Report 2022 – Building a nature-positive society. WWF.

Yates, K. L., et al. (2018). Outstanding challenges in the transferability of ecological models. Trends in ecology & evolution, 33(10), 790–802. 10.1016/j.tree.2018.08.001

