## Supporting Information for "Transferability of stream benthic macroinvertebrate distribution models to drought-related conditions"

^#^Sonja C. Jähnig and Yusdiel Torres-Cambas contributed equally to this work

* Corresponding author:

Graciela Medina-Madariaga

Macroinvertebrate presence and absence records were processed from abundance data obtained from biomonitoring performed by regional authorities and the Senckenberg Institute spanning the years 2007 to 2021. The samples were collected during two seasons, with tributary and low-order streams sampled during spring, and the sites along the main channel of the Kinzig sampled during summer. Macroinvertebrate sampling was performed following the European Water Framework Directive (EU WFD) compliant German national standard sampling protocol (Meier et al. 2006). The field sampling procedure involved using a hand net with a mesh size of 500 μm to collect 20 samples of 25x25 cm squares at each river sampling site, totaling an area of 1.25 square metres of river bottom. Individual specimens, especially those of rare or sensitive species like mayflies and stoneflies, were picked out and preserved in 96% ethanol after being emptied onto a photography tray. Following sample collection, ethanol was exchanged and adjusted for preservation, and samples were stored until processing. In the laboratory, samples were spread evenly on a grid and sorted into coarse and fine fractions using sieve cascades. The coarse fraction, retained on a 2 mm sieve, was further analysed to generate a species list and calculate taxa abundance by classifying them according to the operational taxa list (Haase et al. 2006). Additional sorting was conducted if necessary to ensure accurate abundance estimation.

1.2. Detailed variable processing

The extraction of environmental variables was conducted using R (version 4.2.3; R Core Team, 2023) and QGIS (version 3.28.7; QGIs.org, 2023). To produce bioclimatic variables, rasters downloaded from the German Weather Service (DWD, 2021) were utilised. These files included monthly averages of temperature minima, temperature maxima, and precipitation data at a resolution of 1x1 km. Bioclimatic variables were generated by extracting values from these rasters for the 12 months prior to the sampling date of each occurrence record for every species. The creation of bioclimatic variables for each data point was facilitated by the `biovars()` function from the *dismo* package (Hijmans et al., 2022). Elevation data for each data point was acquired utilising the `sample()` function from the *raster* package (Hijmans & van Etten, 2012) on a 50 m resolution elevation raster, resampled from 1 m LiDAR data obtained from the Hessian Administration for Soil Management and Geoinformation (GDS, 2022) and missing data outside the state of Hesse (<1% of the area) were filled with data from Hawker et al. (2022). Land use maps from InVeKos (2023) and CORINE (2023) were utilised and classified for the primary land coverage types identified (Agriculture, Forest, Urban, Pasture and others) in 2016. The year 2016 was chosen as it represented the last year in the training dataset, and changes in land use during the previously surveyed years were minimal. The upstream contribution of all types of land cover to each data point was computed using the `calculate_exact_attributes` function from the openSTARS package (Kattwinkel et al., 2020), with resulting contributions included in the predictor dataset for each data point. We computed 155 IHAs for each data point using the *EFlowStats* (Thompson et al., 2013) package. Daily flow data for the previous 12 months to sampling were obtained from a SWAT+ model produced for this catchment. The SWAT+ model was calibrated and validated at eight gauges within the catchment and reached good performance scores of the Kling-Gupta-Efficiency (Kling et al. 2012), ranging between 0.72-0.83 (median of 0.77) for the complete simulation period (2006-2023). IHAs were calculated based on the seasonality of sampling. For the occurrence points sampled in spring, IHAs were computed using the previous 12 months counting from April backwards, while for those sampled in summer, the calculation considered the previous 12 months counting from June backwards.

As a result of the data processing, accounting for all categories, 155 predictor variables (five for land use, 19 for bioclimatic variables, elevation, 130 for IHAs; Table S1) were produced for each data point. To streamline this abundance of information, we initiated a rigorous variable selection process for each species before including them into our models. First, for variables in the bioclimatic and land use categories we used the function ‘corSelect’ from the package *fuzzySim* (Barbosa, 2015) to obtain a correlation matrix for all variables, when a pair of variables obtained a correlation higher than 0.7 (Dormann et al., 2013), we retained the one with the lowest variable inflation factor (VIF). IHAs were initially separated into five subcategories (magnitude, duration, frequency, timing and rate; Olden & Poff, 2003), subsequently the same previously mentioned variable selection process was performed for each category. The variable reduction resulted in 33 predictor variables. Second, we used the function ‘select07’ from the package *mecofun* (Zurell, 2023) to select the most influential variables discerning the presence or absence of each taxon using Period1 (drought-free). In order to identify the most simple and flexible models (Yates et al., 2018), for each species the number of variables was selected corresponding to their prevalence in Period1, which according to the rule of thumb equates to one predictor for every 10 occurrences approximately (Harrel et al., 1996; Breiner et al., 2015). Final variable selection was based on the predictors with the lowest Akaike Information Criterion (AIC). The chosen variables were then integrated as predictors in the subsequent SDM fitting process (Table S2).

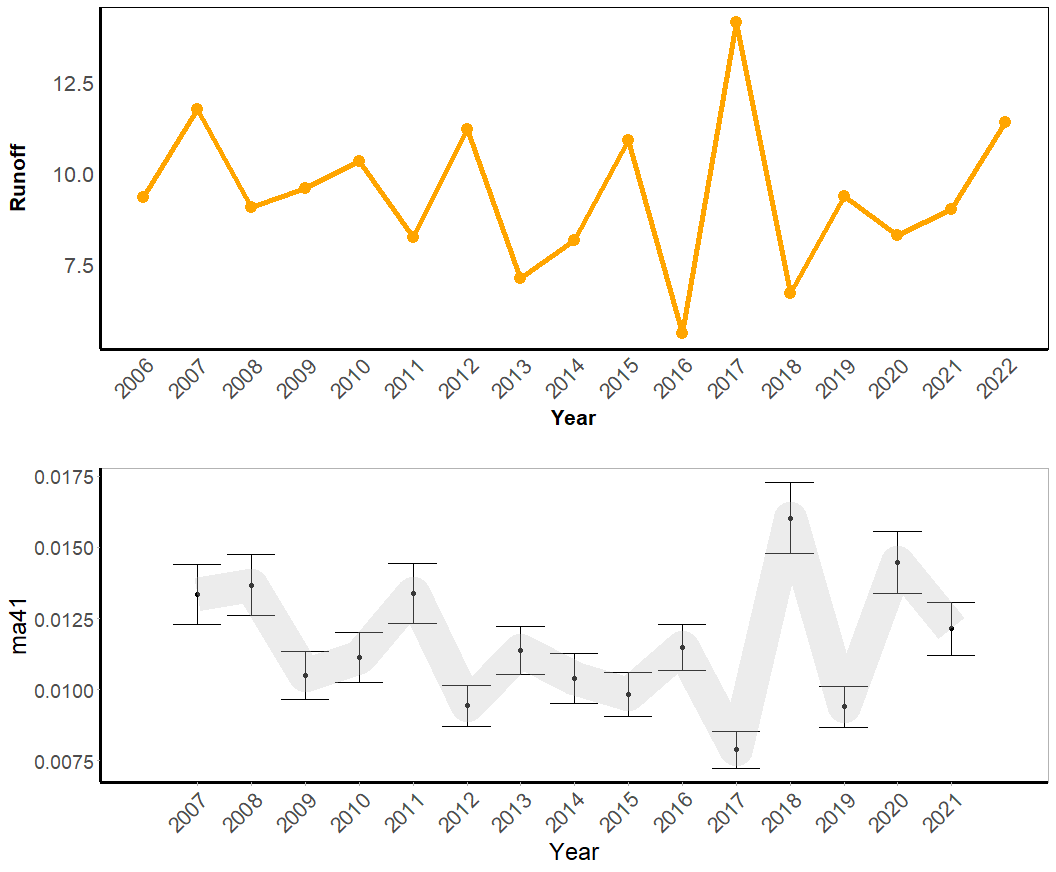

Figure S1. a) 12-month moving average daily streamflow (m³/s) in the Kinzig catchment at outlet (Hanau), during the years 2005-2021. Note that 2016 shows the record lowest runoff in the area. b) Runoff rate (m^3^s^-1^km^2^) calculated across all sampling sites (Fig. 1; using data of the previous 12 months before sampling).

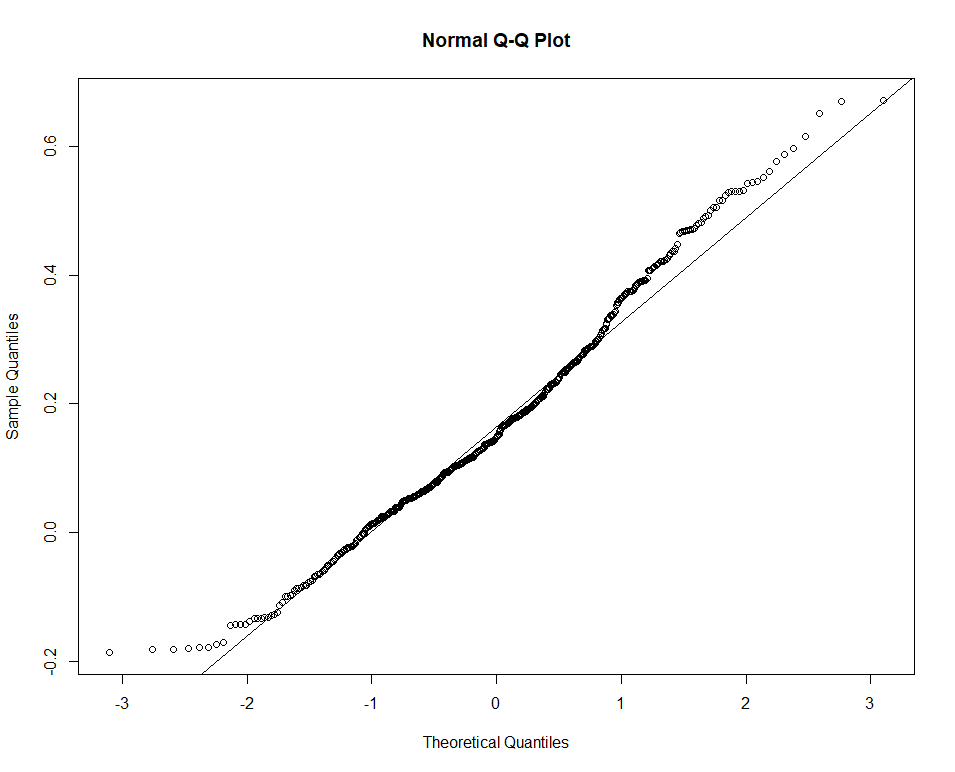

Figure S2. Quantile-quantile plot illustrating the distribution of the response variable AUCgap. The observed skewness in the plot suggests departure from a normal distribution, indicating potential non-normal behaviour in the data. Shapiro-Wilk normality test confirmed abnormality (W = 0.9842, p-value <0.001).

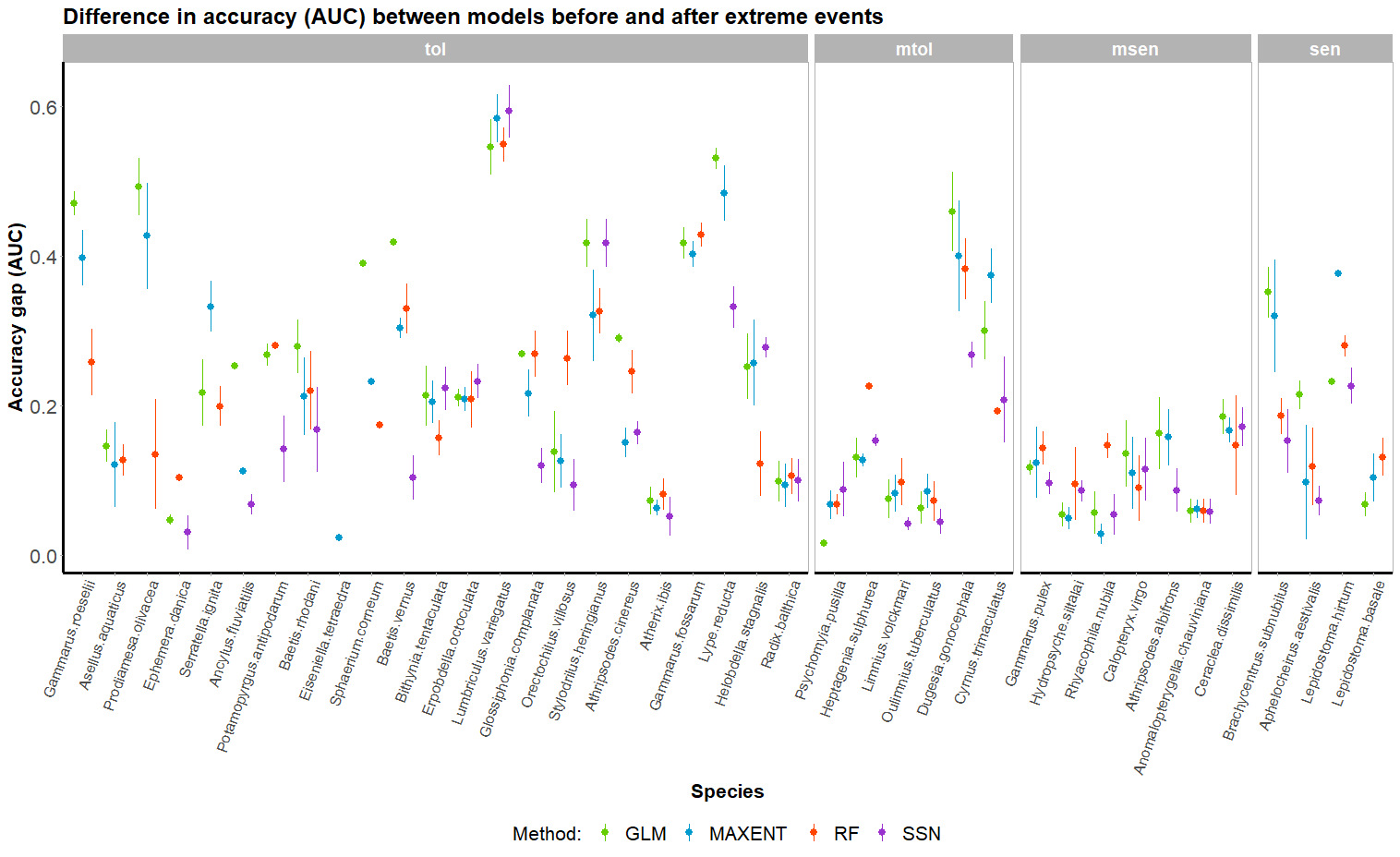

Figure S3. Visual inspection of the effect of species modelled on the AUCgap of models under drought-free and drought-influenced conditions. Species are organised from left to right in a descending number of presences in the dataset, within their tolerance category.

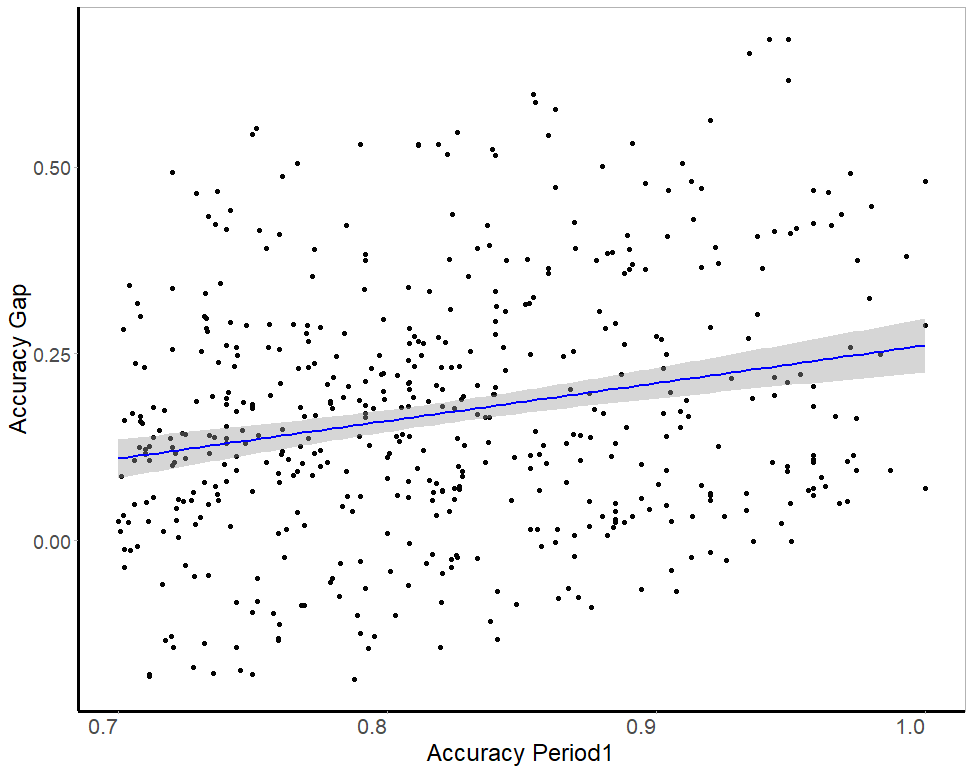

Figure S4. Relationship between the accuracy of models trained and evaluated on Period1 and the accuracy gap (R²= 0.051, p<0.001)

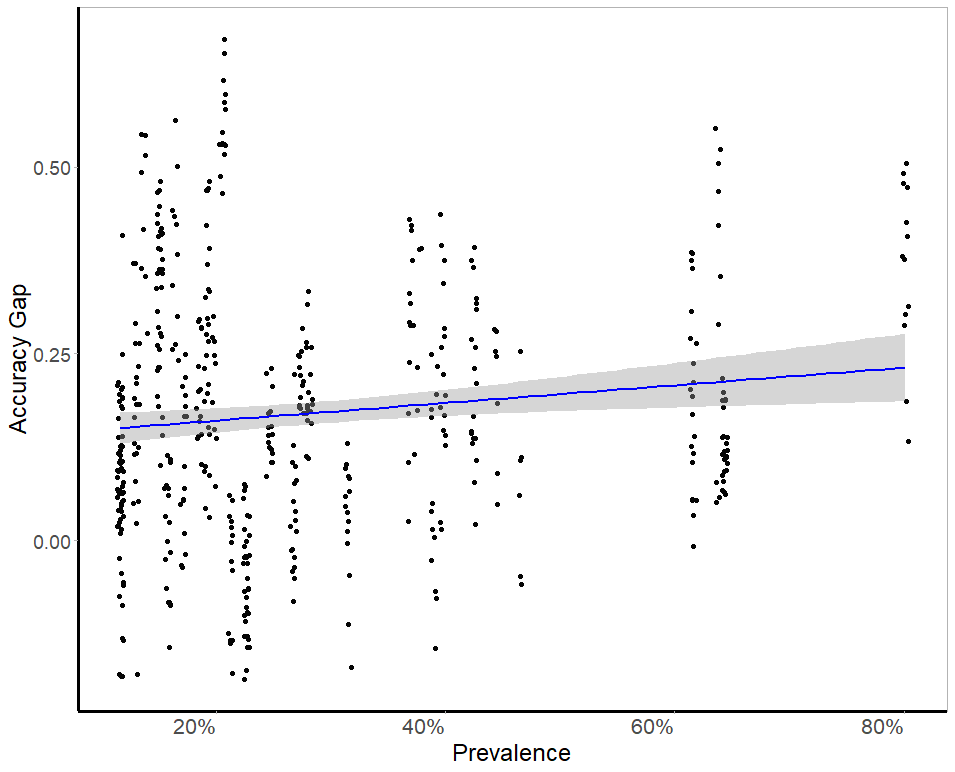

Figure S5. Relationship between the accuracy of models trained and evaluated on Period1 and the accuracy gap (R²= 0.013, p = 0.005)

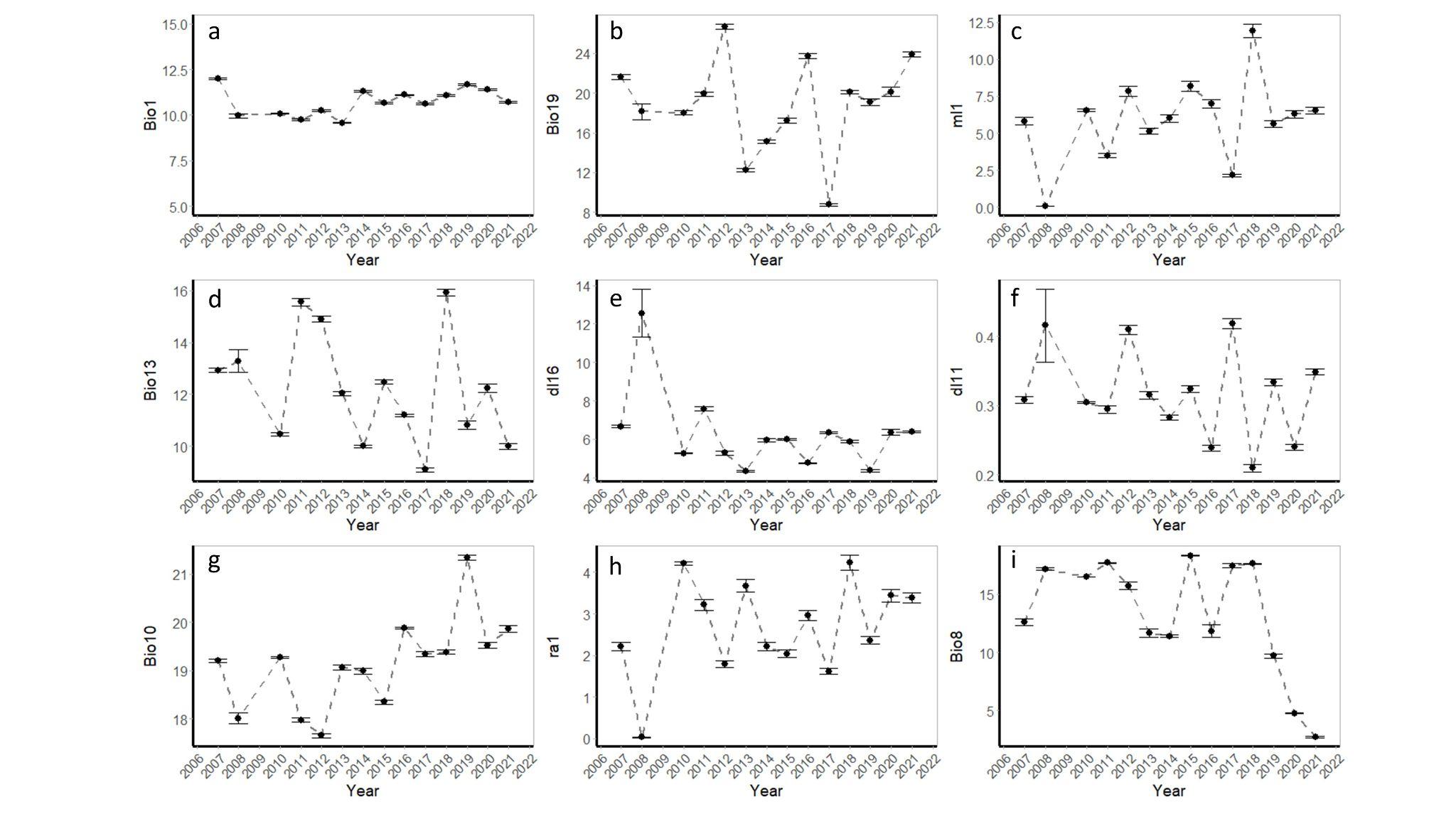

Figure S6. Temporal variability of nine most selected variables (excluding stationary variables such as elevation and land use). Dots represent the mean across sites. Year represent sampling dates and variable values were obtained by using the data of the 12 previous months before sampling dates.

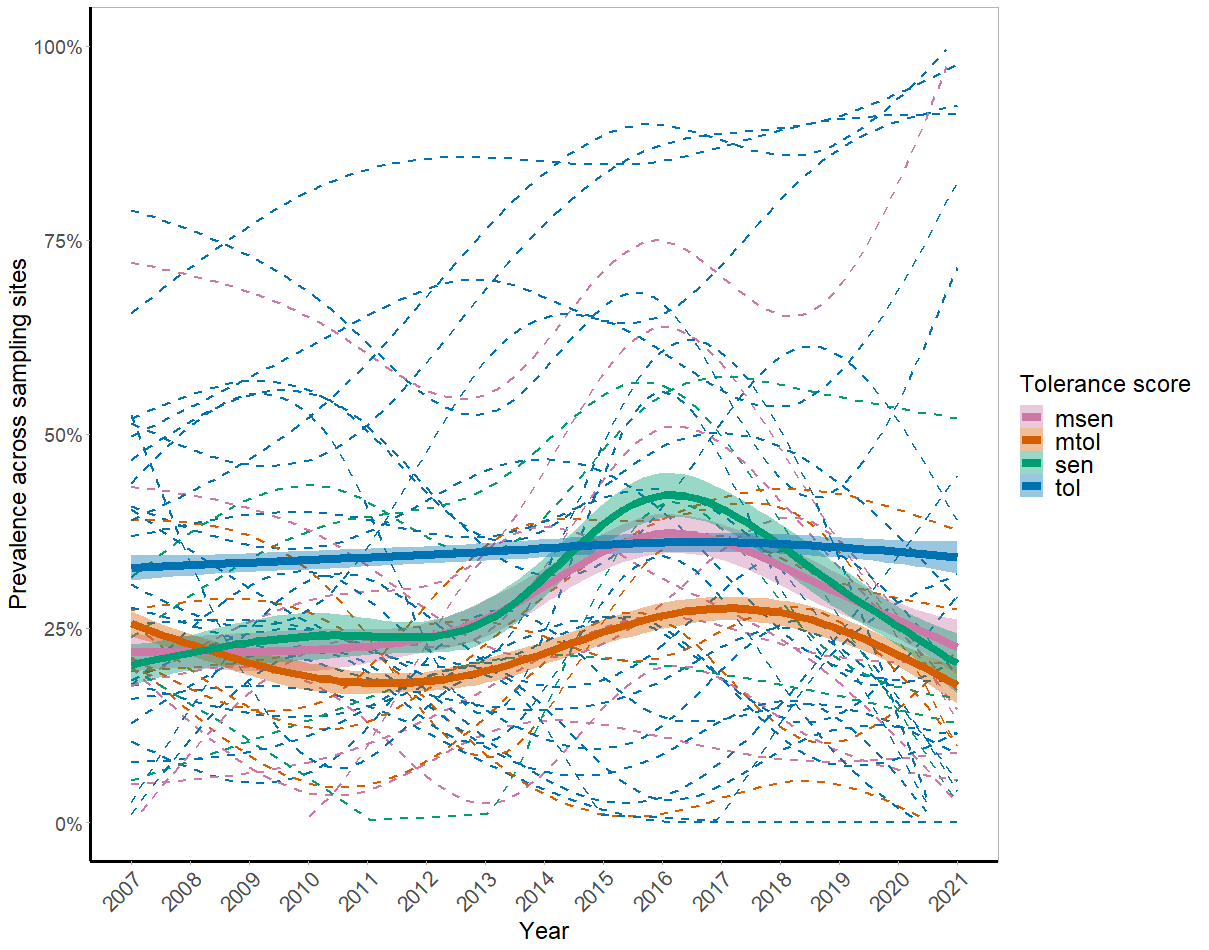

Figure S7. Prevalence of species across sites during the period 2007-2021 according to tolerance score (smoothed lines obtained with GAMs; formula = Prevalence~s(Year, k=6, by=tolerance)). Dashed lines represent individual species coloured by their tolerance score. Thick lines represent the modelled mean prevalence according to tolerance scores and shading represents standard errors.

Table S1. Short description of variables included in the variable selection process (followed by SDM fitting). Additional information on IHA metrics can be found in Olden & Poff (2003).

| Category | Variable | Short description |
| --- | --- | --- |
| Bioclimatic | Bio1 | Annual Mean Temperature |
|  | Bio2 | Mean Diurnal Range (Mean of monthly (max temp - min temp)) |
|  | Bio3 | Isothermality (BIO2/BIO7) (×100) |
|  | Bio4 | Temperature Seasonality (standard deviation ×100) |
|  | Bio5 | Max Temperature of Warmest Month |
|  | Bio6 | Min Temperature of Coldest Month |
|  | Bio7 | Temperature Annual Range (BIO5-BIO6) |
|  | Bio8 | Mean Temperature of Wettest Quarter |
|  | Bio9 | Mean Temperature of Driest Quarter |
|  | Bio10 | Mean Temperature of Warmest Quarter |
|  | Bio11 | Mean Temperature of Coldest Quarter |
|  | Bio12 | Annual Precipitation |
|  | Bio13 | Precipitation of Wettest Month |
|  | Bio14 | Precipitation of Driest Month |
|  | Bio15 | Precipitation Seasonality (Coefficient of Variation) |
|  | Bio16 | Precipitation of Wettest Quarter |
|  | Bio17 | Precipitation of Driest Quarter |
|  | Bio18 | Precipitation of Warmest Quarter |
|  | Bio19 | Precipitation of Coldest Quarter |
| IHAs | dh1 | Annual maxima of 1 day means of daily discharge |
|  | dh2 | Annual maxima of 3 day means of daily discharge |
|  | dh3 | Annual maxima of 7 day means of daily discharge |
|  | dh4 | Annual maxima of 30 day means of daily discharge |
|  | dh5 | Annual maxima of 90 day means of daily discharge |
|  | dh11 | Means of 1 day maxima of daily discharge |
|  | dh12 | Means of 7 day maxima of daily discharge |
|  | dh13 | Means of 30 day maxima of daily discharge |
|  | dh14 | Flood duration 1 |
|  | dh15 | High flow pulse duration |
|  | dh17 | High flow duration 1 |
|  | dh18 | High flow duration 1 |
|  | dh19 | High flow duration 1 |
|  | dh20 | High flow duration 2 |
|  | dh21 | High flow duration 2 |
|  | dl1 | Annual minima of 1 day means of daily discharge |
|  | dl2 | Annual minima of 3 day means of daily discharge |
|  | dl3 | Annual minima of 7 day means of daily discharge |
|  | dl4 | Annual minima of 30 day means of daily discharge |
|  | dl5 | Annual minima of 90 day means of daily discharge |
|  | dl11 | Means of 1 day minima of daily discharge |
|  | dl12 | Means of 7 day minima of daily discharge |
|  | dl13 | Means of 30 day minima of daily discharge |
|  | dl14 | Low exceedance flows |
|  | dl15 | Low exceedance flows |
|  | dl16 | Low flow pulse duration |
|  | dl18 | Number of zero-flow days |
|  | fh1 | High flood pulse count 1 |
|  | fh3 | High flood pulse count 2 |
|  | fh4 | High flood pulse count 2 |
|  | fh5 | Flood frequency 1 |
|  | fh6 | Flood frequency 1 |
|  | fh7 | Flood frequency 1 |
|  | fh8 | Flood frequency 2 |
|  | fh9 | Flood frequency 2 |
|  | fh10 | Flood frequency 3 |
|  | fl1 | Low flood pulse count |
|  | fl3 | Frequency of low flow spells |
|  | ma1 | Mean daily flows |
|  | ma2 | Median daily flows |
|  | ma3 | Variability in daily flows 1 |
|  | ma5 | Skewness in daily flows |
|  | ma7 | Ranges in daily flows |
|  | ma8 | Ranges in daily flows |
|  | ma10 | Spreads in daily flows |
|  | ma11 | Spreads in daily flows |
|  | ma12 | Mean monthly flows |
|  | ma13 | Mean monthly flows |
|  | ma14 | Mean monthly flows |
|  | ma15 | Mean monthly flows |
|  | ma16 | Mean monthly flows |
|  | ma17 | Mean monthly flows |
|  | ma18 | Mean monthly flows |
|  | ma19 | Mean monthly flows |
|  | ma20 | Mean monthly flows |
|  | ma21 | Mean monthly flows |
|  | ma22 | Mean monthly flows |
|  | ma23 | Mean monthly flows |
|  | ma24 | Mean monthly flows |
|  | ma25 | Variability in monthly flows |
|  | ma26 | Variability in monthly flows |
|  | ma27 | Variability in monthly flows |
|  | ma28 | Variability in monthly flows |
|  | ma29 | Variability in monthly flows |
|  | ma30 | Variability in monthly flows |
|  | ma31 | Variability in monthly flows |
|  | ma32 | Variability in monthly flows |
|  | ma33 | Variability in monthly flows |
|  | ma34 | Variability in monthly flows |
|  | ma35 | Variability in monthly flows |
|  | ma36 | Variability across monthly flows 1 |
|  | ma37 | Variability across monthly flows 1 |
|  | ma38 | Variability across monthly flows 1 |
|  | ma39 | Variability across monthly flows 2 |
|  | ma40 | Variability across monthly flows 2 |
|  | ma41 | Mean annual runoff |
|  | mh1 | Mean maximum monthly flows |
|  | mh2 | Mean maximum monthly flows |
|  | mh3 | Mean maximum monthly flows |
|  | mh4 | Mean maximum monthly flows |
|  | mh5 | Mean maximum monthly flows |
|  | mh6 | Mean maximum monthly flows |
|  | mh7 | Mean maximum monthly flows |
|  | mh8 | Mean maximum monthly flows |
|  | mh9 | Mean maximum monthly flows |
|  | mh10 | Mean maximum monthly flows |
|  | mh11 | Mean maximum monthly flows |
|  | mh12 | Mean maximum monthly flows |
|  | mh13 | Variability across maximum monthly flows |
|  | mh14 | Median of annual maximum flows |
|  | mh15 | High flow discharge |
|  | mh16 | High flow discharge |
|  | mh17 | High flow discharge |
|  | mh20 | Specific mean annual maximum flows |
|  | mh21 | High flow volume |
|  | mh22 | High flow volume |
|  | mh24 | High peak flow 1 |
|  | mh25 | High peak flow 1 |
|  | mh27 | High peak flow 2 |
|  | ml1 | Mean minimum monthly flows |
|  | ml2 | Mean minimum monthly flows |
|  | ml3 | Mean minimum monthly flows |
|  | ml4 | Mean minimum monthly flows |
|  | ml5 | Mean minimum monthly flows |
|  | ml6 | Mean minimum monthly flows |
|  | ml7 | Mean minimum monthly flows |
|  | ml8 | Mean minimum monthly flows |
|  | ml9 | Mean minimum monthly flows |
|  | ml10 | Mean minimum monthly flows |
|  | ml11 | Mean minimum monthly flows |
|  | ml12 | Mean minimum monthly flows |
|  | ml13 | Variability across minimum monthly |
|  | ml14 | Mean of annual minimum flows |
|  | ml15 | Low flow index |
|  | ml16 | Median of annual minimum flows 2 |
|  | ml17 | Baseflow index 1 |
|  | ml19 | Baseflow index 2 |
|  | ml20 | Baseflow index 3 |
|  | ml21 | Variability across annual minimum |
|  | ml22 | Specific mean annual minimum flows |
|  | ra1 | Rise rate |
|  | ra2 | Variability in rise rate |
|  | ra3 | Fall rate |
|  | ra4 | Variability in fall rate |
|  | ra5 | No day rises |
|  | ra6 | Change of flow |
|  | ra7 | Change of flow |
|  | ra8 | Reversals |
|  | ra9 | Variability in reversals |
|  | ta1 | Constancy |
|  | th1 | Julian date of annual maximum |
|  | tl1 | Julian date of annual minimum |

Table S2. Species selected to produce SDMs, the variables used to model them, the number of records in the dataset before drought was identified (Period1), and their tolerance score. The names in the column species are in random order.

| Species | Formula | Records | Tolerance |
| --- | --- | --- | --- |
| *Baetis vernus* | fh6 + LU2 + bio10 + Elev + ma34 + bio1 + 1 | 54 | tol |
| *Potamopyrgus antipodarum* | ma25 + ml22 + bio17 + Elev + fl1 + ma35 + bio19 + 1 | 65 | tol |
| *Prodiamesa olivacea* | LU3 + bio19 + bio13 + mh7 + fh1 + dl16 + LU4 + fl3 + ta1 + LU1 + 1 | 93 | tol |
| *Brachycentrus subnubilus* | mh11 + dh4 + ma11 + ma33 + fl1 + bio13 + 1 | 58 | sen |
| *Cyrnus trimaculatus* | ra4 + ma29 + bio1 + 1 | 22 | mtol |
| *Asellus aquaticus* | LU3 + dl11 + fl3 + ma32 + ml22 + bio1 + Elev + bio2 + ml1 + LU4 + 1 | 94 | tol |
| *Gammarus pulex* | Elev + bio10 + bio1 + dh4 + dl11 + mh11 + ra4 + bio2 + ml22 + dl16 + 1 | 94 | msen |
| *Gammarus roeselii* | LU4 + LU2 + LU1 + fl3 + ta1 + dl16 + fh7 + bio17 + bio7 + ma30 + ma32 + ma41 + 1 | 117 | tol |
| *Serratella ignita* | LU1 + ml1 + LU2 + LU3 + fl3 + ma33 + ra1 + ma30 + fl1 + 1 | 90 | tol |
| *Anabolia nervosa* | ma28 + LU1 + th1 + bio8 + ra8 + 1 | 47 | msen |
| *Calopteryx virgo* | ma25 + ml22 + ra1 + 1 | 25 | msen |
| *Bithynia tentaculata* | ml1 + ra1 + dl16 + ma11 + bio8 + 1 | 41 | tol |
| *Sphaerium corneum* | dh4 + dl11 + ml11 + fl3 + bio1 + ma32 + 1 | 55 | tol |
| *Rhyacophila nubila* | bio1 + tl1 + LU1 + ml22 + 1 | 31 | msen |
| *Aphelocheirus aestivalis* | dh4 + mh11 + ma11 + dl11 + dl16 + mh5 + 1 | 57 | sen |
| *Lepidostoma hirtum* | dl16 + bio13 + bio19 + 1 | 22 | sen |
| *Psychomyia pusilla* | ma11 + LU1 + LU2 + fh4 + 1 | 39 | mtol |
| *Athripsodes albifrons* | ma35 + ma31 + 1 | 19 | msen |
| *Ancylus fluviatilis* | bio19 + Elev + dh15 + bio10 + bio13 + LU2 + mh12 + 1 | 68 | tol |
| *Gammarus fossarum* | mh7 + ml1 + bio19 + 1 | 22 | tol |
| *Ephemera danica* | Elev + LU2 + ta1 + bio1 + bio8 + ml20 + bio10 + ma41 + dl11 + 1 | 90 | tol |
| *Heptagenia sulphurea* | bio1 + ra1 + ma32 + ml1 + 1 | 36 | mtol |
| *Athripsodes cinereus* | ml1 + th1 + mh7 + 1 | 27 | tol |
| *Baetis rhodani* | ml1 + ra1 + bio19 + bio13 + bio8 + dl16 + bio1 + 1 | 62 | tol |
| *Lumbriculus variegatus* | ma33 + ma26 + ra6 + 1 | 30 | tol |
| *Dendrocoelum lacteum* | LU2 + Elev + 1 | 16 | tol |
| *Erpobdella octoculata* | mh12 + LU2 + mh5 + mh11 + 1 | 40 | tol |
| *Lype reducta* | ma28 + ma33 + 1 | 20 | tol |
| *Eiseniella tetraedra* | ra7 + ma34 + dl11 + fh6 + ra6 + dh15 + 1 | 58 | tol |
| *Hydropsyche siltalai* | Elev + fl1 + mh7 + bio19 + ml1 + 1 | 46 | msen |
| *Dugesia gonocephala* | bio19 + bio13 + dh15 + 1 | 24 | mtol |
| *Stylodrilus heringianus* | LU2 + mh5 + bio1 + 1 | 28 | tol |
| *Glossiphonia complanata* | ml1 + ra1 + dl16 + 1 | 29 | tol |
| *Helobdella stagnalis* | bio7 + LU4 + 1 | 19 | tol |
| *Radix balthica* | LU1 + bio10 + 1 | 17 | tol |
| *Atherix ibis* | LU1 + bio10 + LU4 + 1 | 23 | tol |
| *Oulimnius tuberculatus* | LU4 + mh12 + LU1 + bio8 + 1 | 33 | mtol |
| *Limnius volckmari* | mh12 + dl11 + Elev + bio13 + 1 | 33 | mtol |
| *Orectochilus villosus* | bio13 + bio19 + ma41 + 1 | 28 | tol |
| *Anomalopterygella chauviniana* | Elev + dh4 + 1 | 17 | msen |
| *Lepidostoma basale* | mh1 + ml11 + 1 | 17 | sen |
| *Ceraclea dissimilis* | th1 + bio2 + 1 | 17 | msen |

Table S3. Kruskal-Wallis tests for groups. The table presents the results of Kruskal-Wallis tests for different groups, with respect to various response variables. The response variables include: Period1Acc: AUC of models trained and evaluated on Period1. Period2Acc: AUC of models trained and evaluated in Period1 and validated on Period2. AUCgap: The difference between Period1Acc and Period2Acc. Each response variable is analysed with respect to the group variable explaining it. Degrees of freedom (df) and p-values (p) from the Kruskal-Wallis tests are reported.

| Response Variable | Group variable | Chi-squared | df | p |
| --- | --- | --- | --- | --- |
| Period1Acc | Method | 8.7736 | 3 | 0.03 |
| Period1Acc | Tolerance score | 6.3929 | 3 | 0.093 |
| Period2Acc | Method | 23.619 | 3 | <0.001 |
| Period2Acc | Tolerance score | 110.17 | 3 | <0.001 |
| AUCgap | Method | 10.537 | 3 | 0.014 |
| AUCgap | Tolerance score | 103.0538 | 3 | <0.001 |

Table S4. Anova type III test results for our full model (formula 2). Chisq values obtained through Type III Wald chi square tests, degrees of freedom (DF), p values represent significance of the variable, values under 0.1 in italics and values under 0.05 are in bold and italics.

|  | Chisq | DF | P value |
| --- | --- | --- | --- |
| ***Method*** | ***28.818*** | ***3*** | ***<0.001*** |
| ***Tolerance*** | ***8.267*** | ***3*** | ***0.040*** |
| ***Method:Tolerance*** | ***18.613*** | ***9*** | ***0.028*** |

References

Barbosa, A. M. (2015). fuzzySim: applying fuzzy logic to binary similarity indices in ecology. Methods in Ecology and Evolution, 6(7), 853-858. <https://doi.org/10.1111/2041-210X.12372>

Breiner, F. T., Guisan, A., Bergamini, A., & Nobis, M. P. (2015). Overcoming limitations of modelling rare species by using ensembles of small models. Methods in Ecology and Evolution, 6(10), 1210-1218.

Copernicus Land Monitoring Service 2024. European Environment Agency (EEA) (2021). <https://land.copernicus.eu/pan-european/corine-land-cover>. Assessed 25 March 2024.

Dormann, C. F., et al. (2013). Collinearity: a review of methods to deal with it and a simulation study evaluating their performance. Ecography, 36(1), 27-46.

DWD. (2021) German Weather Service climate data. <https://opendata.dwd.de/climate_environment/CDC/observations_germany/climate/>

Friedman, J., Hastie, T., & Tibshirani, R. (2010). Regularization paths for generalized linear models via coordinate descent. Journal of statistical software, 33(1), 1.

GDS. (2022). 1m Digital Elevation Model. Hessian Administration for Soil Management and Geoinformation. Available online: <https://gds.hessen.de/INTERSHOP/web/WFS/HLBG-Geodaten-Site/de_DE/-/EUR/ViewDownloadcenter-Start>

Haase, P., Lohse, S., Pauls, S., Schindehütte, K., Sundermann, A., Rolauffs, P., Hering, D. (2004), Assessing streams in Germany with benthic invertebrates: development of a practical standardised protocol for macroinvertebrate sampling and sorting. Limnologica 34:349–365.

<https://doi.org/10.1016/S0075-9511(04)80005-7>

Haase, P., Sundermann, A., Schindehütte, K. (2006) Informationstext zur Operationellen Taxaliste als Mindestanforderung an die Bestimmung von Makrozoobenthosproben aus Fließgewässern zur Umsetzung der EU-Wasserrahmenrichtlinie in Deutschland. <https://gewaesser-bewertung-berechnung.de/files/downloads/perlodes/Operationelle_Taxaliste_Begleittext.pdf>.

Harrell, F. E., Lee, K. L., & Mark, D. B. (1996). Multivariable prognostic models: issues in developing models, evaluating assumptions and adequacy, and measuring and reducing errors. Statistics in medicine, 15(4), 361-387.

Hawker, L., Uhe, P., Paulo, L., Sosa, J., Savage, J., Sampson, C., & Neal, J. (2022). A 30 m global map of elevation with forests and buildings removed. Environmental Research Letters, 17(2), 024016.

Hijmans, R.J, Phillips, S., Leathwick, J. and Elith, J. (2022), Package ‘dismo’. Available online at: <http://cran.r-project.org/web/packages/dismo/index.html>.

Hijmans, R.J & van Etten, J. (2012). raster: Geographic analysis and modeling with raster data. R package version 2.0-12. [http://CRAN.R-project.org/package=raster](http://cran.r-project.org/package=raster)

InVeKos. (2021). Crop data based on the German agricultural subsidies program. <https://www.zi-daten.de>. Assessed 25 March 2024.

Kattwinkel, M., Szöcs, E., Peterson, E., & Schäfer, R. B. (2020). Preparing GIS data for analysis of stream monitoring data: The R package openSTARS. Plos one, 15(9), e0239237.

Kling, H., Fuchs, M., Paulin, M., 2012. Runoff conditions in the upper Danube basin under an ensemble of climate change scenarios. Journal of Hydrology. 424–425, 264–277. <https://doi.org/10.1016/j.jhydrol.2012.01.011>

Meier, C., Haase, P., Rolauffs, P., Schindehütte, K., Schöll, F., Sundermann, A., & Hering, D. (2006). Methodisches Handbuch Fließgewässerbewertung - Handbuch zur Untersuchung und Bewertung von Fließgewässern auf der Basis des Makrozoobenthos vor dem Hintergrund der EG-Wasserrahmenrichtlinie (Vol. Stand: Mai 2006). <https://gewaesser-bewertung-berechnung.de/>

Nguyen, H. H., et al. (2024). Stream macroinvertebrate communities in restored and impacted catchments respond differently to climate, land-use, and runoff over a decade. Science of The Total Environment, 929, 172659. <https://doi.org/10.1016/j.scitotenv.2024.172659>

Olden, J.D. and Poff, N.L. (2003). Redundancy and the choice of hydrologic indices for characterizing streamflow regimes. River Research and Applications, 19, 101-121. <https://doi.org/10.1002/rra.700>

QGIS.org (2023). QGIS Geographic Information System. QGIS Association. <http://www.qgis.org>

R Core Team (2023). R: A language and environment for statistical computing. R Foundation for Statistical Computing, Vienna, Austria. [https://www.R-project.org/](https://www.r-project.org/).

Thompson, J., Archfield, S. A., Kennen, J., & Kiang, J. (2013). EflowStats: An R package to compute ecologically-relevant streamflow statistics. In AGU Fall Meeting Abstracts (Vol. 2013, pp. H43E-1508).

Yates, K. L., et al. (2018). Outstanding challenges in the transferability of ecological models. Trends in ecology & evolution, 33(10), 790-802.

Zurell, D. (2023). mecofun: useful functions for macroecology and species distribution modelling. version 0.1.2. University of Potsdam, Potsdam. <https://gitup.uni-potsdam.de/macroecology/mecofun>
